# CYB5R4 sustains endothelial proliferation and ischemia-induced angiogenesis by maintaining RRM2-dependent nucleotide balance

**DOI:** 10.1101/2025.09.17.676962

**Authors:** Shuai Yuan, Scott A. Hahn, Nicole C. Colussi, Steven J. Mullett, Stacy L. Gelhaus, Francisco J. Schopfer, Adam C. Straub

## Abstract

Angiogenesis is essential for revascularization in peripheral artery disease (PAD), yet pro-angiogenic therapies remain inconsistent. Here, we identify a cytosolic reductase, CYB5R4, as an intrinsic regulator of endothelial proliferation and ischemia-induced angiogenesis. In mice, global haploinsufficiency or inducible endothelial deletion of *Cyb5r4* delayed perfusion recovery after femoral artery ligation and reduced capillary expansion. CYB5R4 is known to promote stearoyl-CoA desaturase (SCD) activity and is required for the *de novo* synthesis of monounsaturated fatty acids. In human endothelial cells, *CYB5R4* silencing impaired proliferation with G1-S arrest that was not rescued by monounsaturated fatty acids and differed from the loss of SCD, indicating an SCD-independent mechanism. RNA sequencing with Bayesian network inference highlighted the ribonucleotide-reductase subunit RRM2 as a key downstream mediator. RRM2 overexpression partially restored proliferation. Integrated untargeted metabolomics and targeted nucleotide quantification revealed an imbalanced nucleotide pool in CYB5R4-deficient cells. These findings support a model in which CYB5R4 sustains endothelial proliferation and ischemia-driven angiogenesis by maintaining RRM2-dependent nucleotide balance. Targeting the CYB5R4-RRM2 axis may improve therapeutic angiogenesis in PAD.

## Introduction

Peripheral artery disease (PAD) is a common atherosclerotic disease characterized by narrowed limb arteries and reduced blood flow, affecting over 200 million people worldwide^1,2^. The most severe manifestation is chronic limb ischemia (CLI), presenting ischemic rest pain, non-healing wounds, and gangrene, often necessitating surgical revascularization or even amputation^3^. While surgical or endovascular revascularization is the standard of care, a substantial subset of PAD patients are poor candidates for these procedures^4,5^. In ischemic tissues, the body attempts to compensate by activating angiogenesis, the sprouting of new capillaries from existing vessels, via hypoxia-inducible mediators such as vascular endothelial growth factor (VEGF). Intuitively, therapeutic angiogenesis, or augmenting new vessel growth in ischemic tissue, has been explored for decades as a strategy to improve perfusion in PAD^6^. However, despite encouraging preclinical studies, clinical trials delivering pro-angiogenic factors, such as VEGF, in PAD patients have shown little to moderate benefit, especially for limb salvage and functional mobility^7–9^. These disappointments highlight the need for a deeper understanding of the biological limitations and alternative approaches to promote effective angiogenesis in PAD.

Endothelial cells (ECs) are central mediators of angiogenesis, requiring coordinated proliferation and migration in response to ischemia. These processes demand metabolic reprogramming: while glycolysis provides the majority of ATP, fatty acid β-oxidation supplies carbon to the tricarboxylic acid cycle for aspartate production, a precursor for de novo nucleotide biosynthesis^10,11^. Inhibition of this pathway depletes nucleotide pools and impairs EC proliferation and sprouting, and metabolic derangements such as diabetes similarly disrupt angiogenesis in PAD^12^. Thus, EC metabolic fitness has emerged as a key determinant of angiogenic capacity.

Angiogenesis is shaped by a redox balance. Cardiometabolic risk factors, including diabetes, smoking, and aging, elevate oxidative stress, reduce nitric oxide bioavailability, and blunt neovascularization^13^. Yet reactive oxygen species also function as essential second messengers in growth factor signaling, as exemplified by NADPH oxidase 4 (NOX4)–derived hydrogen peroxide (H_2_O_2_), which promotes angiogenic signaling^14^. This duality underscores the need for precise enzymatic control of redox balance in ECs. Members of the cytochrome b5 reductase (CYB5R1-5) family are increasingly recognized as such regulators^15^. For example, CYB5R3 was recently shown to cooperate with NOX4 to facilitate EC H_2_O_2_ production^16^. On the other hand, CYB5R4 is structurally distinct, harboring both a cytochrome b5–like domain and a reductase domain that has been shown to regulate fatty acid monodesaturation through stearoyl-CoA desaturase^17–19^. Prior work has shown that loss of CYB5R4 in mice disrupts unsaturated fatty acid synthesis, induces lipotoxic stress, and sensitizes cells to oxidative injury, highlighting its essential role in metabolic and redox regulation^20,21^. Thus, the intricate interplay between pro-angiogenic redox cues and oxidative damage emphasizes redox-regulating enzymes as critical mediators of EC homeostasis.

These observations led us to hypothesize that angiogenic ECs, which require robust lipid synthesis and nucleotide production, may also depend on CYB5R4 activity. To investigate this, we employed a causal inference approach using Bayesian network analysis, a probabilistic graphical model that can infer putative cause-and-effect relationships from transcriptomic data^22–24^. By integrating RNA sequencing with Bayesian inference, we aimed to identify CYB5R4-dependent regulatory programs that may influence EC proliferation and angiogenesis. Our findings uncover a previously unrecognized regulatory paradigm in ECs and vascular homeostasis, whereby CYB5R4 orchestrates metabolic–redox control through its functional interplay with the ribonucleotide reductase regulatory subunit M2 (RRM2).

## Methods

### Public genetics and expression resources

CYB5R4 constraint metrics were queried from gnomAD using the official API (https://gnomad.broadinstitute.org/api)^25^. Expression of CYB5R4 across arteries and muscle was assessed from the GTEx portal (dbGaP Accession phs000424.v8.p2). GWAS summary statistics were obtained from the Common Metabolic Diseases Knowledge Portal (CMDKP) [cmdkp.org *CYB5R4* Region page. 2025 July 17 https://hugeamp.org/region.html?chr=6&end=84777142&phenotype=PAD&start=84469375(RRID:SCR_020937)]^26^. Gene traits were aligned to Hg19 reference genome for visualization. The traits shared by the GTEx and CMDKP were used in a linear regression analysis.

### Mouse strains, allele engineering, and genotyping

A Cyb5r4 null allele and a floxed allele were generated on the C57BL/6J background using CRISPR-Cas9 genome editing^27^. Single guide RNAs, targeting *Cyb5r4* Exon 3 (**Supplementary Table 1**), were synthesized by PCR as previously described^28^. Founders, carrying the null or floxed allele, were backcrossed for 4 generations and inbred for 2 generations before experimental use (**Fig. 1c,g**). Endothelium-specific deletion was achieved by crossing mice carrying the floxed allele with Cdh5(PAC)-CreERT2 mice^16,29^. Tamoxifen (40 mg/kg daily for 10 days) was used to induce Cre activity in 8-week-old mice. Mice were housed under a 12-h light/dark cycle with ad libitum access to chow and water. All animal studies were approved by and in compliance with the University of Pittsburgh Institutional Animal Care and Use Committee.

**Fig. 1:**
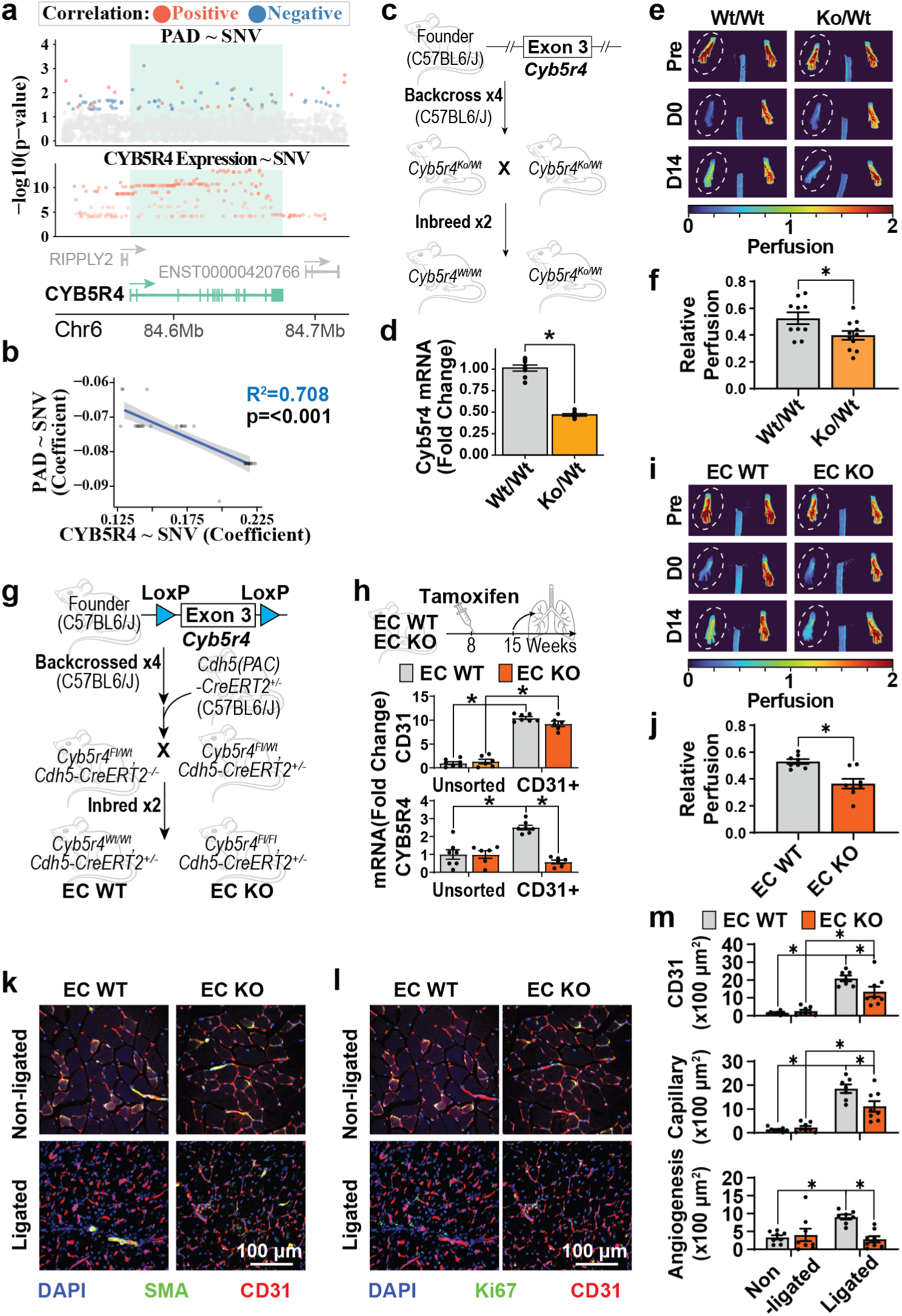
CYB5R4 expression promotes ischemia-induced angiogenesis. **(a)** The dot plot shows the association between single-nucleotide variants (SNV) of *CYB5R4* and peripheral artery disease or *CYB5R4* expression. Statistics are retrieved from the Common Metabolic Diseases Knowledge Portal and the Genotype-Tissue Expression Portal. Significant associations (p < 0.05) are shown as red (positive) or blue (negative). **(b)** A linear regression analysis shows a negative correlation between a variant’s coefficients for PAD and *CYB5R4* expression. **(c, g)** Illustrations of the generation and breeding of the global *Cyb5r4* knockout and tamoxifen-inducible endothelium-specific *Cyb5r4* knockout mice. **(d)** *Cyb5r4* mRNA expression in lungs isolated from the global *Cyb5r4* knockout strain. N= 8; * indicates p < 0.05 using a Student’s t-test. (**e, i**) Representative laser speckle contrast images (LSCI) before (Pre) and after (D0) femoral artery ligation, and 14 days after the surgery (D14). **(f, j)** Quantification of the LSCI results 14 days after femoral artery ligation. * indicates p < 0.05 comparing the Cyb5r4 homozygous wildtype (Wt/Wt) and heterozygous knockout (Ko/Wt) or the endothelium-specific inducible knockout (EC KO) group and the WT control (EC WT) groups using Student’s t-tests. N = 10 (Wt/Wt), 11 (Ko/Wt), 8 (EC WT), and 8 (EC KO) **(h)** Cyb5r4 mRNA expression in unsorted and endothelial (CD31+) fractions of lung cells. * indicates p < 0.05 between linked groups using Two-way ANOVA and Tukey’s post hoc test. N = 7 (EC WT) and 6 (EC KO) **(k, l)** Representative images of immunostaining for α smooth muscle actin (SMA), PECAM-1 (CD31), and Ki67 in mouse gastrocnemius muscle sections. **(m)** Quantifications of CD31, capillary, and angiogenesis areas. N=8; * indicates p < 0.05 between linked groups using two-way ANOVA and Tukey’s post hoc test.

### Viability, body weight, and glucose tolerance

Mendelian ratios were assessed at weaning by genotyping the entire litter. Body weight was recorded before weaning (14-28 days postnatal). Only the weight on Day 21 was reported. In case Day 21 was missing, the data was imputed by linear regression using the two nearest measurements. For glucose tolerance testing, mice were fasted for 16 hours and injected intraperitoneally with D-glucose (2 g/kg). Tail vein glucose was measured at 0, 15, 30, 60, 90, and 120 min using a handheld glucometer (Zoetis Diagnostics, AlphaTrak3).

### Femoral artery ligation

At 8 weeks of age, mice from the global knockout strain (Wt/Wt and Ko/Wt, **Fig. 1c**) and the endothelium-specific knock strain (EC WT and EC KO, **Fig. 1g**) were injected with tamoxifen (33 mg/kg) intraperitoneally for 10 consecutive days to induce CYB5R4 knockout in the endothelium. At 15 weeks of age, unilateral hindlimb ischemia was induced as described previously with minor modifications^30^. Under isoflurane anesthesia (1.5-2.0% in oxygen) and aseptic technique, a skin incision was made in the left groin to expose the femoral neurovascular bundle. The left femoral artery was ligated at the branch points of the lateral circumflex femoral artery and the popliteal artery^31^. The intervening arterial segment was excised, while the femoral vein and nerve were preserved. Buprenorphine (3.25 mg/kg, subcutaneously) was administered for analgesia, and mice were placed on a warming pad until recovery.

### Laser speckle contrast imaging (LSCI)

Blood perfusion in both hindlimbs was measured before ligation, immediately after ligation, and 14 days after ligation using the PeriCam PSI NR laser speckle contrast imager using the PIMSoft PSI software. Mice were anesthetized with 1-1.5% isoflurane and positioned prone on a heated stage to maintain 37 °C. Regions of interest (ROIs) were defined over the plantar surfaces of both paws. Measurements were collected at 25 frames per second and averaged every second until a stable perfusion rate was observed. Perfusion was averaged over the ROI within a two-minute measurement window using the PIMSoft PSI software. The reported perfusion values were normalized to contralateral non-ischemic limbs for each animal. Mice with less than 70% blood flow reduction immediately after ligation were excluded from the study. For visualization, binary image files containing variance and intensity data were exported from PIMSoft PSI software to produce a temporally averaged image in Python, following the instrument’s manual.

### Skeletal muscle processing and immunofluorescence

Gastrocnemius muscles were harvested 14 days after ligation, fixed in 4% paraformaldehyde, cryoprotected in 30% sucrose, embedded in OCT, and cryosectioned (10 μm). Sections were permeabilized with 0.5% Triton X-100, blocked in 10% normal serum, and incubated with primary antibodies against CD31 (R&D/Novus, AF3628, 1:100), FITC-conjugated α-smooth muscle actin (Sigma, F3777, 1:500), and Ki67 (Abcam, ab16667). Fluorophore-conjugated secondary antibodies (Alexa Fluor 594/647, Invitrogen) were used for detection, and nuclei were counterstained with DAPI. Images were acquired on a Nikon A1 confocal microscope using identical settings across conditions. Endothelium, capillary, and angiogenesis areas were defined and quantified using binary thresholds. Both quantification and visualization were performed using Python.

### Endothelial cell culture and treatments

Primary human aortic endothelial cells (HAECs) (Lonza, CC-2535) were cultured in endothelial growth medium (Lonza, CC-3156) supplemented with growth factors and antibiotics (Lonza, CC-4176) at 37 °C with 5% CO₂. Cells with population doubling times less than 12 were used in this study. For gene silencing, cells were transfected with siRNAs targeting CYB5R4 (Invitrogen, Silencer Select, s27597 or s27596), SCD (Invitrogen, Silencer Select, s12503), or a non-targeting control (Invitrogen, 4390843) using Lipofectamine 3000 (Invitrogen, L3000008) according to the manufacturer’s instructions. Briefly, HAECs were seeded in 12-well plates at a density of 20,000-23,000 cells/cm^2^. Cells were allowed to settle overnight until they were treated with a liposome complex containing 1.5 μl Lipofectamine 3000 and siRNA to reach 10 nM or otherwise stated final concentration. After overnight incubation, cells were washed and supplied with fresh cell culture media. In fatty acid (FA) supplementation experiments, cells were treated with bovine serum albumin (BSA)-conjugated FAs (FA:BSA at 5:1 molar ratio) 48 hours after media change. Unless noted otherwise, cells were collected 72 hours after media change.

### Immunoblotting

For immunoblotting, cells were lysed directly into 2× Laemmli sample buffer containing 5% β-mercaptoethanol and denatured at 95 °C for 10 min. Equal protein amounts were separated on 4-12% Bis-Tris NuPAGE gels and transferred to nitrocellulose membranes. Membranes were blocked for 1 h at room temperature in PBS with 3% BSA and 0.05% Tween-20, then incubated overnight at 4 °C with primary antibodies to CYB5R4 (Santa Cruz, sc-390569, 1:1,000), SCD (Cell Signaling, 2438, 1:1,000), RRM2 (Proteintech, 11661-1-AP, 1:1,000), and α-tubulin (Sigma, T6074, 1:10,000). After washing, IRDye 680RD or 800CW secondaries (LI-COR, 1:10,000) were applied, and signals were captured on a LI-COR imager. Densitometry was performed with LI-COR Image Studio; bands were normalized to α-tubulin and expressed as fold change relative to controls.

### Quantitative PCR

Tissue or cells were lysed in the RLT buffer for RNA isolation using the RNeasy Mini Kit (Qiagen, 74104). The cDNA library was synthesized using SuperScript IV VILO (Invitrogen, 11756050). Diluted cDNA samples and primer sets spanning at least one intron of the target gene were used to set up the quantitative PCR (qPCR) reactions with Power SYBR Green PCR Master Mix (Applied Biosystems, 4367659). The PCR was run on a QuantStudio 5 platform in the 384-well format. The Ct values were used to calculate the expression difference between the gene of interest and the housekeeping gene (ΔCt) (*RPS13* for human and *Gapdh* for mouse) (**Supplementary Table 1**). The fold changes of experimental groups over the control group (ΔΔCt) were used for statistics.

### Endothelial cell isolation

Mouse lungs were isolated without perfusion. Minced lung tissues were digested in 1 mg/ml collagenase A (Roche, 10103578001) in phosphate-buffered saline (PBS, calcium and magnesium-free) at 37 °C for 1 hour. Cells were dislodged by passing through an 18-G needle and filtered through a 70 μm mesh. Cells were washed and suspended in PBS with 2mM EDTA, 4.5mg/mL D-glucose. For each sample, ECs were labeled with 2 μl CD31 antibody (BD Biosciences, 550274) at 4 °C for 30 minutes and immobilized by anti-rat IgG Dynabeads (Invitrogen, 11036D) at 4 °C for 30 minutes. The bead-bound ECs were then washed and captured on a magnet. The EC and unsorted cell fractions were lysed directly for qPCR.

### EdU incorporation and cell-cycle analysis

48 hours after siRNA transfection and media change, HAECs were starved in endothelial basal media for 4 hours and then treated with 10 μM 5’-ethynyl-2’-deoxyuridine (EdU) in fully supplemented media for 16 hours. Cells were trypsinized, fixed with 4% paraformaldehyde, and stained using Click-iT Plus EdU Alexa Fluor 647 Flow Cytometry Kit (Invitrogen, C10635). EdU⁺ fractions were quantified by high-content imaging or flow cytometry.

For cell-cycle profiling, siRNA-transfected HAECs were collected 72 hours after media change. Cells were fixed in 70% ethanol, treated with 100 μg/ml RNase A (NEB, T3018L), stained with 50 μg/ml propidium iodide (Invitrogen, P1304MP), and analyzed by flow cytometry using the cell cycle analysis function in FlowJo v10.8.1.

### LC-MS/MS analysis of fatty acid coenzyme A conjugates

To assess stearoyl-CoA desaturase (SCD) activity, siRNA-transfected HAECs were incubated with deuterated stearic acid (17,17,18,18,18-D_5_, 98%) (Cambridge Isotope Laboratories, DLM-2712-0.1) for 16 h. Cells were collected, and cell pellets were precipitated with 80% Methanol containing 60 pmol of heptadecanoyl CoA. Samples were vortexed, centrifuged at 1,000 x g for 5 min at 4°C. The supernatant containing CoA conjugates was analyzed by HPLC-HR-MS/MS using gradient solvent systems consisting of water containing 0.1% NH_4_OH (solvent A) and acetonitrile containing 0.1% NH_4_OH (solvent B). Samples were resolved for qualitative characterization using a reverse-phase HPLC column (2 × 150 mm, 5 µm Luna C8(2) column; Phenomenex) at a 0.35 ml/min flow rate. Samples were loaded onto the column at 5% B, maintained for 0.3 min, and eluted with a linear increase in solvent B from 5-45% B over 3 min, then 45-65% B over 3 min. The system was re-equilibrated to initial conditions over 4 minutes. The analysis was performed using a QTrap 6500+ triple quadrupole mass spectrometer (Sciex, San Jose, CA) in the negative ion mode. Source temperature was 650 °C, curtain gas: 50, ionization spray voltage: 5500, GS1: 55, GS2: 60. For collision induced fragmentations leading to the specific neutral loss fragment of m/z 507, the following settings were used: declustering potential: 70 V, entrance potential: 10 V, collision energy: 35 V and collision cell exit potential: 8 V. 18:1-d5 and 18:0-d5 peaks were quantified by the area under the curve and expressed as the 18:1-d5/18:0-d5 ratio.

### RNA sequencing

Total RNA was isolated from HAECs transfected with a titration series of CYB5R4 siRNA (Invitrogen, Silencer Select, s27597) and non-targeting controls using RNeasy Mini Kit (Qiagen, 74104). RNA samples were sent to Novogene for unstranded paired-end bulk RNA sequencing on an Illumina platform. Reads were trimmed, aligned to GRCh38 using STAR, and gene-level counts were obtained with featureCounts. Lowly expressed genes were filtered. Differential expression analysis was performed using DESeq2 with default settings by comparing the 10 nM CYB5R4 and non-targeting siRNA-transfected samples. An independent validation dataset was generated using a second, non-overlapping siRNA targeting CYB5R4 (Invitrogen, Silencer Select, s27597) processed identically. Gene set enrichment analysis and pathway overrepresentation tests were performed using ClusterProfiler. Statistics reported by DESeq2 and ClusterProfiler were used for visualization in R.

### Bayesian network inference and hub identification

To infer putative causal structure among highly differentially expressed genes (DEGs, FDR < 10^-10^), we built directed acyclic graph (DAG) models using bnlearn^32^. Structure learning was performed with tabu search using the Bayesian Gaussian score (BGe) on log2-transformed count per million values. The search started with 1000 random graphs using Melancon’s and Philippe’s method. Importantly, we retrieved human protein-protein interaction data from the STRING database with a score threshold of 400. Importantly, in the Castelo and Siebes prior^33^, arcs present in the STRING database were slightly rewarded (+0.1) to aid the network structure learning. Edge confidence was assessed as the frequency of an edge across bootstrap graphs relative to null networks generated by permuting sample labels. A consensus network retained edges passing an FDR threshold determined from the null. Node centrality was computed using igraph, including authority and hub scores from the HITS algorithm, in- and out-degree, and out/in edge ratios. Candidates with high authority score, hub score, and out/in edge ratios were prioritized as hubs.

### Doxycycline-inducible overexpression screen

To test predicted hubs functionally, we created a small cDNA library [*RRM2* (Sino Biological, HG18284-U), *TK1* (Addgene, 100544), *BIRC5* (Addgene, 136348), *AARS1* (Transomic, DQ891634), *CDC6* (Addgene, 109332), *TPX2* (Addgene, 54285), *CYB5R3*^16^, and *EGFP*^34^] in a doxycycline-inducible lentiviral system (Addgene, 171123). The lentiviral backbone was modified to express a blasticidin resistance gene (VectorBuilder). Terminal tags were removed from the open reading frames before they were cloned into the lentiviral backbone using PCR. All cloning processes were verified with Sanger sequencing. Lentiviruses were packaged using the third-generation packing system (Invitrogen), and functional viral particles were quantified using a colony formation assay in HEK293 cells.

HAECs were transduced with 1 multiplicity of infection (MOI) in 6-well plates. Cells were selected in 5 μg/ml blasticidin for 7 days before they were pooled together. Pooled HAECs were transfected with *CYB5R4* siRNA (Invitrogen, Silencer Select, s27597) as described above. Meanwhile, gene expression was induced by the addition of 100 ng/ml doxycycline for 48 hours. Genomic DNA was purified using the DNeasy Blood & Tissue Kit (Qiagen, 69504) and treated with 100 μg/ml RNase A. Transgene copies were quantified using primer sets spanning at least one intron in the target gene (**Supplementary Table 1**), so only the transgene sequence is amplified in qPCR. The viral gene *RRE* was used as the housekeeping control for the total viral load in the ΔCt calculation. The ΔΔCt values for doxycycline-treated samples over vehicle control samples are used as the fold enrichment.

### Targeted RRM2 rescue experiment

The RRM2 cDNA was cloned into a bicistronic EGFP coexpression vector pCIGX^34^. Plasmid transfection was performed as we previously reported^34^. HAECs were plated in 6-well plates at a density of 23,000 cells/cm^2^. Lipofectamine 3000 transfection mixture was set up using 0.2 μg plasmid DNA, 3.75 μl Lipofectamine 3000, and 0.4 μl P3000 reagent according to the manufacturer’s instructions. Cells were incubated with the transfection reagent and 5 μM MRT67301 for 2 hours in fully supplemented growth media. After the incubation, cells were thoroughly washed and then transfected with siRNA as described above overnight in the presence of 5 μM MRT67301. After changing media the next morning, cells were allowed to grow for 48 hours before they were used for the EdU incorporation experiment.

### Proximity labeling (APEX2) and streptavidin pulldown

To map proteins in the immediate vicinity of CYB5R4, a C-terminal APEX2 (Addgene, 85823) fusion of CYB5R4 in the pCIGX vector was expressed in HAECs. The experiment was performed as we previously reported^16^. Briefly, cells were incubated with 500 μM biotin-phenol (Cayman, 26997) for 30 minutes, followed by 60 seconds of exposure to 100 μM H_2_O_2_ to catalyze proximity-dependent biotinylation. Reactions were quenched (10 mM sodium ascorbate, 10 mM sodium azide, 5 mM Trolox [Cayman, 10011659] in phosphate-buffered saline, pH 8), cells were lysed, and biotinylated proteins were enriched with streptavidin-conjugated Dynabeads (Invitrogen, 11205D). After stringent washes, eluates were analyzed by immunoblotting for candidate interactors.

### Proximity ligation assay (PLA)

Endogenous spatial proximity between CYB5R4 and RRM2 was assessed using a Duolink in situ PLA kit. HAECs grown on coverslips were fixed, permeabilized, and incubated with primary antibodies against CYB5R4 (Santa Cruz, sc-390569, 1:50) and RRM2 (Proteintech, 11661-1-AP, 1:50) from different host species. Species-specific PLA probes were applied, followed by ligation and rolling-circle amplification. Amplified signals were visualized as puncta by fluorescence microscopy. Puncta per nucleus were quantified across fields using identical exposure settings and automated image analysis in Python.

### Untargeted metabolomics and joint pathway analysis

For global metabolite profiling, siRNA-transfected HAECs were trypsinized, pelleted, and snap-frozen in liquid nitrogen. Cell pellets were sent to Novogene for the untargeted metabolomics service. Extracts were prepared in methanol:water (4:1, v/v) containing internal standards, subjected to three freeze-thaw cycles (liquid N_2_/dry ice/ice), clarified by centrifugation, and the supernatant was used for LC-MS. Data were acquired on an ExionLC AD coupled to a SCIEX TripleTOF 6600+ using T3 reversed-phase (water/acetonitrile +0.1% formic acid) and BEH Amide HILIC (ammonium formate, pH 10.6) methods in positive and negative electrospray ionization modes (4 µl injection, 0.40 ml/minute, 40 °C). MS1 (50-1000 Da) and MS2 (25-1000 Da) spectra were collected with vendor default source potentials and temperatures. Pooled QC was injected every 10 samples, blanks were interspersed, and internal standards (CV ≤ 15%) monitored run stability.

Original data was converted to mzXML (ProteoWizard), features were detected/aligned in XCMS, peaks with >50% missing per group were excluded, remaining missing values were KNN-filled, and signal drift was corrected by SVR. Compounds were annotated against in-house/public databases and MetDNA; only IDs with a score > 0.7 and QC CV < 0.3 were kept, then positive/negative modes were merged. Intensities were normalized to protein (BCA). Differential metabolites were called using orthogonal partial least squares discriminant analysis (OPLS-DA) variable importance in projection (VIP) > 1 plus p-value < 0.05 (t-test or Wilcoxon test).

Differential metabolites with KEGG annotations and DEGs from the RNA sequencing (FDR < 10^-6^) experiment were submitted to MetaboAnalyst (queried in May 2024) to perform hypergeometric tests on metabolic pathways. P-values were integrated by combining queries. The pathway impact was calculated with degree centrality.

### Targeted nucleotide quantification

Metabolic quenching and nucleotide phosphate metabolite pool extraction were performed by adding ice-cold 2:2:1 (acetonitrile:methanol:water) at a ratio of 500 µL per 10^6^ cells. [^15^N_5_]-adenosine 5’-monophosphate (AMP), [^15^N_5_]-adenosine 5’-diphosphate (ADP), [^15^N_5_]-adenosine 5’-triphosphate (ATP), [^15^N_3_]-cytidine-5’-monophosphate (CMP), [^2^H_14_]-cytidine-5’-triphosphate (CTP), [^15^N_5_]-guanosine-5’monophosphate (GMP), [^15^N_5_]-guanosine-5’-triphosphate (GTP), [^15^N_2_, ^13^C_10_]-thymidine-5’-monophosphate (TMP), [^15^N_2_, ^13^C_10_]-thymidine-5’-triphosphate (TTP), [^15^N_2_]-uridine-5’-monophosphate (UMP), [^15^N_2_]-uridine-5’-triphosphate (UTP) (Sigma-Aldrich) were added to the sample lysates as an internal standards at a final concentration of 10 µM. Samples are homogenized by vortexing for 3 minutes before clearing of protein by centrifugation at 16,000 x g. Cleared supernatant (400 µL) was dried to completion under N_2_ gas and resuspended in 40 µL of lysis buffer and 2 µL was subjected to online LC-MS analysis.

Analyses were performed by untargeted LC-HRMS. Briefly, Samples were injected via a Thermo Vanquish UHPLC and separated over a reversed-phase Thermo HyperCarb porous graphite column (2.1×100 mm, 3 μm particle size) maintained at 55°C. For the 30-minute LC gradient, the mobile phase consisted of the following: solvent A (water/7.5 mM ammonium acetate/0.05% ammonium hydroxide) and solvent B (ACN/0.05% ammonium hydroxide). The gradient was the following: 0-1.8 minutes 5% B, increase to 65% B over 12.2 minutes, continue increasing to 98% B over 0.1 minutes, hold at 98%B for 4 minutes, reequilibrate at 5%B for 12 minutes. The Thermo Exploris 240 mass spectrometer was operated in positive ion mode, scanning in ddMS^2^ mode (2 μscans) from 300 to 800 *m/z* at 120,000 resolution with an AGC target of 2x10^5^ for full scan, 2x10^4^ for MS^2^ scans using HCD fragmentation at stepped 20,35,50 collision energies. Source ionization setting was 3.4 kV spray voltage. Source gas parameters were 50 sheath gas, 10 auxiliary gas at 350 °C, and 1 sweep gas. Calibration was performed prior to analysis using the Pierce^TM^ FlexMix Ion Calibration Solutions (Thermo Fisher Scientific). Integrated peak areas were then extracted manually using Quan Browser (Thermo Fisher Xcalibur ver. 2.7). Relative group differences were reported by normalizing metabolite peak areas to the peak area of the internal standards.

## Statistics and reproducibility

For *in vivo* experiments, each mouse was considered a biological replicate (n=1). Unless specified, *in vitro* experiments were repeated on different days, and each replicate is considered n=1. The exact n values and the statistical test used for each panel are provided in the figure legends. Results were reported as mean ± standard error of the mean (SEM). For two-group comparisons, two-tailed Student’s t-tests were applied after verifying normality and variance homogeneity. Otherwise, non-parametric tests were used, for comparisons with more than two groups or factors, one- or two-way ANOVA with Tukey’s post-hoc test was used. Analyses were performed with GraphPad Prism 10 or customized code in R and Python.

## Data availability

Upon publication, the RNA sequencing data generated from this study will be deposited in Gene Expression Omnibus (GEO). Metabolomics data (raw files and processed matrices), imaging datasets, and other raw data will be available in Figshare. Code used for RNA sequencing data analysis, Bayesian network inference, image analysis, and metabolomics preprocessing will be available in GitHub/Figshare.

## Results

### CYB5R4 expression is negatively associated with PAD

We sought to determine whether CYB5R4 gene expression correlates with peripheral artery disease PAD in the human population. Genome-wide association studies (GWAS) curated by the Common Metabolic Disease Knowledge Portal (CMDKP) did not show a strong genome-wide association of *CYB5R4* single-nucleotide variants (SNVs) and PAD prevalence (**Fig.1a**). However, we found that *CYB5R4* showed a moderate depletion of loss-of-function variants according to the Genome Aggregation Database (gnomAD) (**Supplementary** Fig. 1). Indeed, loss-of-function variants are rare in the GWAS datasets, making it challenging to evaluate whether CYB5R4 protein function affects PAD. Similarly, *CYB5R4* variants curated by the Adult Genotype-Tissue Expression (GTEx) project are all positively associated with *CYB5R4* mRNA expression in artery and muscle tissues. Among variants shared by CMDKP and GTEx, the effects of the variants on CYB5R4 expression were positively associated with their effects on PAD (**Fig. 1b**).

### Global *Cyb5r4* knockout mice exhibit sub-Mendelian viability

To investigate the impact of CYB5R4 deletion in vivo, we created global *Cyb5r4* knockout mice and endothelium-specific tamoxifen-inducible knockout mice by targeting exon 3 of the gene (**Fig. 1c, 1g**). Our global *Cyb5r4* knockout mice showed decreased body weight at weaning (21 days) (**Supplementary** Fig. 2a) and hyperglycemia in the glucose tolerance test at 12 weeks of age (**Supplementary** Fig. 2g) as reported in an independent mouse strain^20,35^. Similar to the loss-of-function constraint in humans, the *Cyb5r4* knockout mouse strain is sub-viable, with fewer homozygotes (*Ko/Ko*) than predicted by Mendelian ratios (**Supplementary** Fig. 2b). In comparison, the floxed homozygotes (*Fl/Fl*) showed normal body weight and Mendelian ratios (**Supplementary** Fig. 2e**, 2f**). To rule out the possibility of off-target CRISPR/Cas9 editing, we crossed the floxed mice with a global Cre recombinase expresser (*CMV-Cre*). The resultant knockout strain also showed decreased body weight and viability (**Supplementary** Fig. 2c**, 2d**).

### CYB5R4 deficiency impairs ischemia-induced vascular remodeling in mice

Global heterozygous *Cyb5r4* knockout mice (*Ko/Wt*) showed 50% lower *Cyb5r4* mRNA expression (**Fig. 1d**) compared to the littermate wildtype control mice (*Wt/Wt*), while exhibiting normal body weight (**Supplementary** Fig. 2a**, 2c**), viability (**Supplementary** Fig. 2b**, 2d**), and blood glucose level (**Supplementary** Fig. 2g). We performed unilateral femoral artery ligation on the heterozygous knockout mice to model the impact of decreased *CYB5R4* expression on PAD. We quantified the relative blood perfusion in the hindlimbs using laser speckle contrast imaging (LSCI) before and after the ligation (**Fig.1e**). The blood flow recovery was delayed in the *Ko/Wt* mice compared to the *Wt/Wt* control mice (**Fig. 1f**).

We then used the endothelium-specific inducible Cyb5r4 knockout mice (EC KO) to examine whether endothelium CYB5R4 is required for angiogenesis under ischemia. Tamoxifen injection efficiently decreased CYB5R4 expression specifically in ECs isolated from the EC KO lungs compared to the littermate wildtype control (EC WT) (**Fig. 1h**). The loss of CYB5R4 in the ECs was sufficient to cause impaired blood flow recovery in the hindlimb after femoral artery ligation (**Fig. 1i, 1j**). Two weeks after the ligation, the gastrocnemius muscle was isolated from both the ligated and non-ligated limbs. We performed immunostaining for CD31, SMA, and Ki67 for ECs, smooth muscle cells, and proliferating cells (**Fig. 1k, 1l**). The quantification showed that the endothelium (CD31^+^), capillary (CD31^+^SMA^-^), and angiogenesis (Ki67^+^ capillary) areas are increased in the ligated muscle, which was dampened in EC KO compared to EC WT (**Fig. 1m**).

### CYB5R4 promotes EC proliferation independent of fatty acid monodesaturation

Silencing of *CYB5R4* in human aortic endothelial cells (HAECs) using siRNA showed a decreased wound recovery rate compared to the non-targeting siRNA control (siNT) in a scratch assay (**Fig. 2a-b**). CYB5R4 is required by stearyl-CoA desaturase (SCD) for the *de novo* synthesis of monounsaturated fatty acids in the mouse liver^20^, and the loss of SCD activity causes cell cycle arrest^36^. We confirmed that the loss of CYB5R4 in HAECs inhibited the SCD activity, as measured by the lower conversion of deuterated stearic acid-*d5* (18:0-*d5*) into oleic acid-*d5* (18:1-*d5*); silencing *SCD* (siSCD) further decreased the 18:1-*d5* /18:0-*d5* ratio (**Fig. 1c**). SCD silencing also inhibited scratch wound recovery (**Fig. 2d, 2e**). However, further analyses indicate that CYB5R4 and SCD may regulate cell proliferation via different mechanisms.

**Fig. 2:**
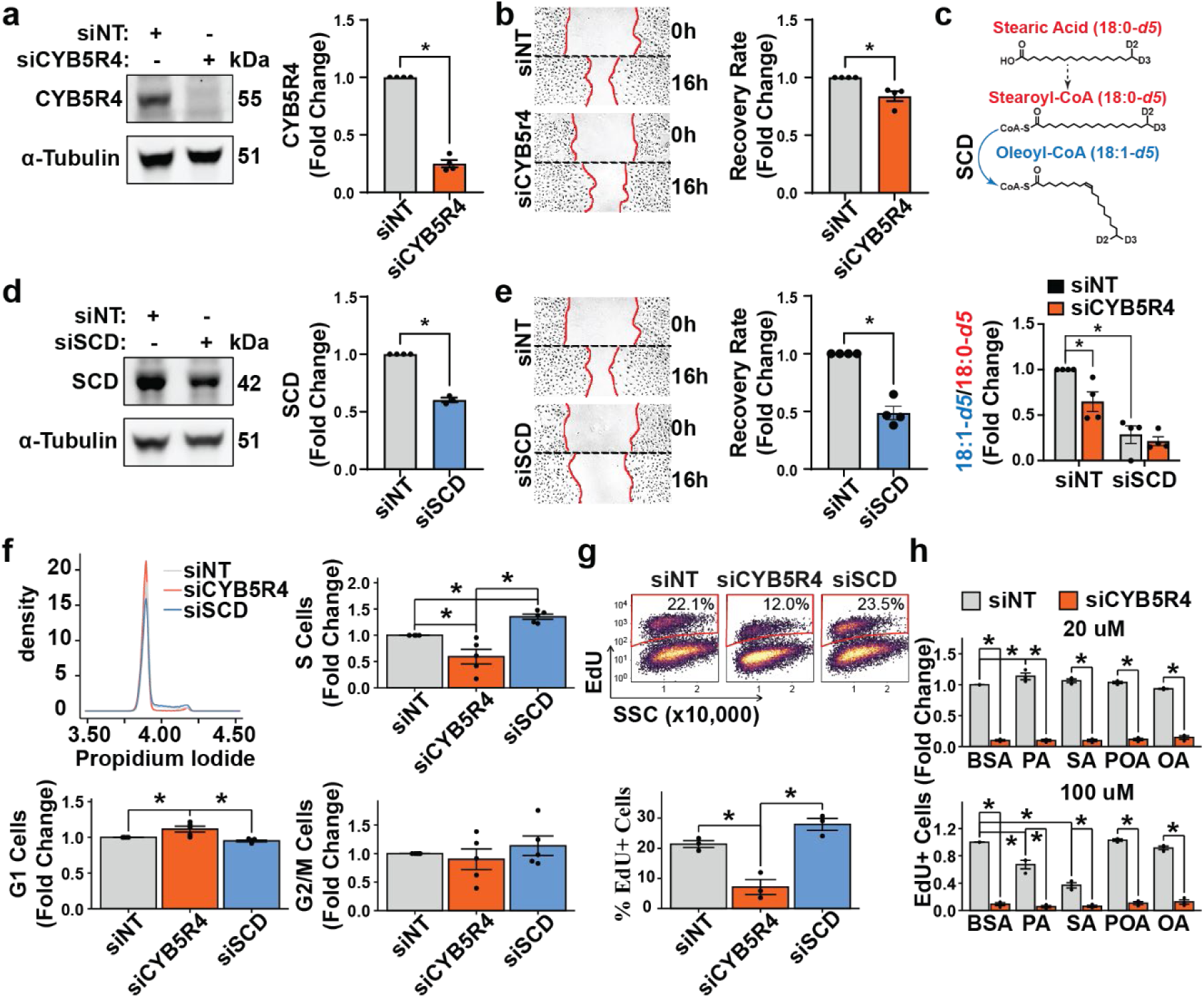
Effects of CYB5R4 and SCD on endothelial proliferation. HAECs were transfected with non-targeting (siNT), *CYB5R4*-targeting (siCYB5R4), or *SCD*-targeting (siSCD) siRNA. **(a, c)** Gene silencing efficiency was verified using immunoblotting. N = 4; * indicates p < 0.05 using Student’s t-tests. **(b, d)** After siRNA transfection, cells were allowed to grow to confluence for 48 hours and used in a scratch wound healing assay. The representative images show the wound edges (highlighted in red) at the beginning and the end of the assay. The recovered wound areas are quantified in the right panels. N = 4; * indicates p < 0.05 using Student’s t-tests. **(e)** HAECs were treated with deuterium (D2)-labeled stearic acid (18:0) to measure the conversion rate of stearic acid to oleic acid (18:1). N = 4; * indicates p < 0.05 between linked groups using two-way ANOVA and Tukey’s post hoc test. **(f)** The representative density plot and quantification of cell cycle analysis using siRNA-transfected HAECs. N = 5; * indicates p < 0.05 between linked groups using one-way ANOVA and Tukey’s post-hoc test. **(g)** The representative dot density plot showing the gating strategy and the quantification of 5-ethynyl-2’-deoxyuridine (EdU) using siRNA-transfected HAECs. N = 3; * indicates p < 0.05 between linked groups using one-way ANOVA and Tukey’s post-hoc test. **(h)** After siRNA transfection, HAECs were treated with palmitic (PA), stearic (SA), palmitoleic (POA), or oleic acids (OA) and used in the EdU incorporation assay. N = 3; * indicates p < 0.05 between linked groups using one-way ANOVA and Tukey’s post-hoc test.

Cell cycle analysis showed that siCYB5R4 decreased the S-phase population while increasing the G1 population, suggesting that the G1-to-S transition is compromised by *CYB5R4* silencing (**Fig. 1f**). In comparison, the loss of SCD increased S-phase cells (**Fig. 1f**). This discrepancy in S-phase cells between siCYB5R4 and siSCD was confirmed by labeling S-phase HAECs with 5-ethynyl-2’-deoxyuridine (EdU) (**Fig. 1g**). Furthermore, supplementation of monounsaturated fatty acids, such as palmitoleic and oleic acids, did not increase the S-phase population in siCYB5R4 HAECs, indicating that CYB5R4 regulates EC proliferation in an SCD-independent manner.

### CYB5R4 regulates genes involved in angiogenesis-related pathways

To discover novel regulatory mechanisms of CYB5R4 in ECs, we performed RNA sequencing in *CYB5R4*-silenced HAECs. The gene set enrichment analysis indicated that CYB5R4 regulated pathways essential for angiogenesis, such as cell cycle, DNA replication, and regulation of actin cytoskeleton (**Fig. 3a**). Biosynthesis of unsaturated fatty acids and fatty acid metabolism were also affected by *CYB5R4* knockdown. Notably, core enrichment genes involved in the aforementioned pathways were suppressed by *CYB5R4* silencing, suggesting a potential downregulation of these pathways. In comparison, core enrichment related to reactive oxygen species increased in *CYB5R4* knockdown cells. The expression of individual core enrichment genes can be found in **Supplementary** Fig. 3.

**Fig. 3:**
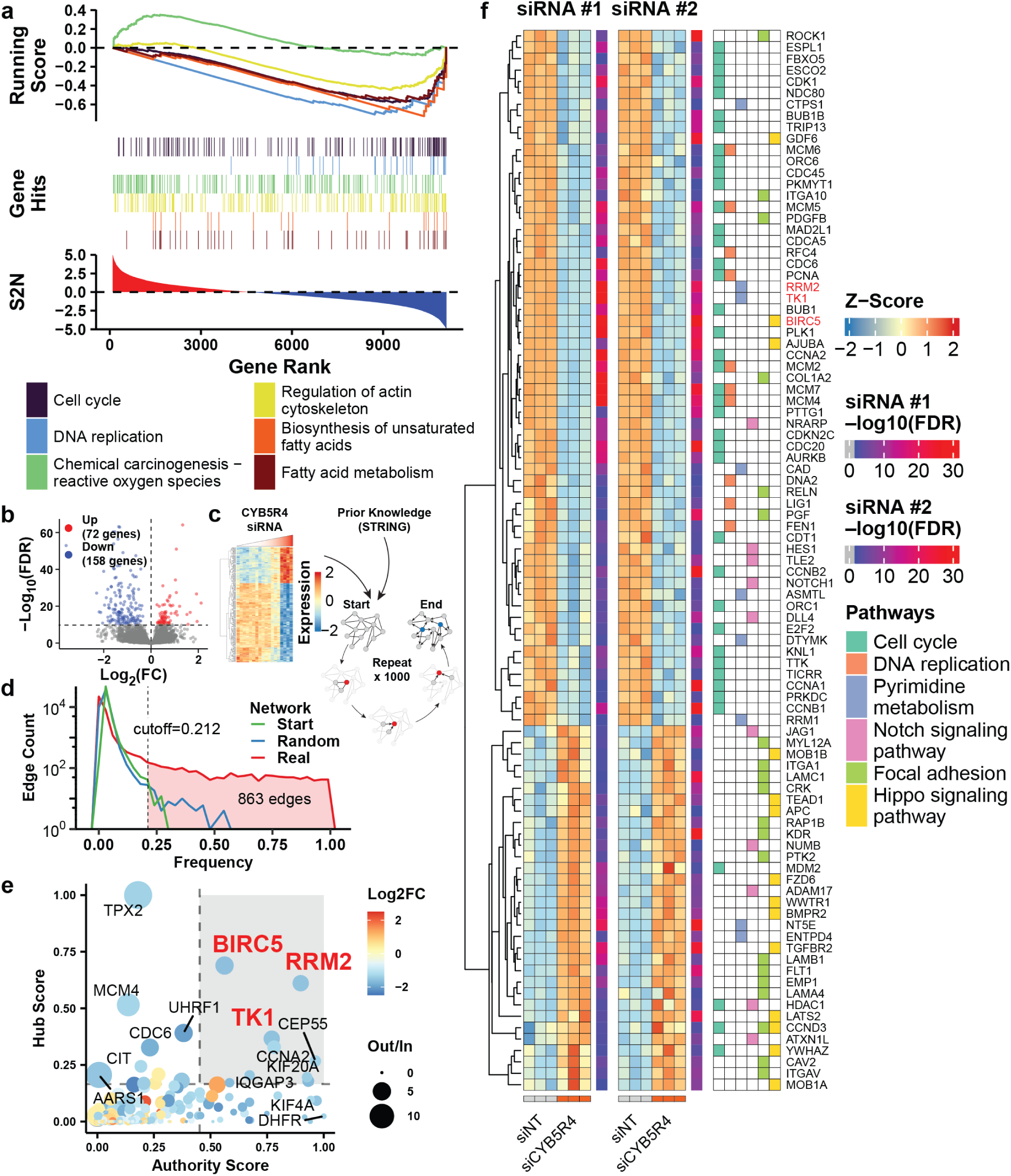
Causal inference with RNA sequencing data. HAECs ere transfected with 10 μM non-targeting (siNT) or various concentrations (0.04-10 μM) of *CYB5R4*-targeting (siCYB5R4) siRNA for RNA sequencing. **(a)** The three horizontally aligned panels show selected significant (FDR < 0.05) Kyoto Encyclopedia of Genes and Genomes (KEGG) pathways in a gene set enrichment analysis. The bottom panel shows the signal-to-noise ratio (S2N) of the 10 μM siCYB5R4 group over 10 μM siNT, which is used to rank genes in descending order. The middle panel shows the rank of genes annotated for selected KEGG pathways, while the top panel shows the running score in the gene set enrichment analysis. **(b)** The volcano plot shows the log2-transformed fold change [log2(FC)] and log10-transformed false discovery rate [log10(FDR)] of differentially expressed genes (DEGs) (10 μM siCYB5R4 over 10 μM siNT). **(c)** The strategy for causal inference using Bayesian networks. The heatmap shows gene-wise scaled and centered expression (z-score) in HAECs treated with different concentrations of siCBY5R4. The expression values of significant DEGs shown in **(b)** (FDR < 10^-10^) are used to update a gene regulatory network based on the STING database. The network structure is learned via changing its edges from a random start point and optimizing the Bayesian information criterion (BIC) score. The process is repeated to generate 1000 different networks. **(e)** The dot plot shows important network metrics used to determine central hub genes. The coordinates of a dot are Kleinberg’s hub and authority centrality scores. The size of a dot represents the ratio of outward edge count to inward edge count. The color of a dot shows the log2(FC) of 10 μM siCYB5R4 over 10 μM siNT. Central hub genes are highlighted in red. **(f)** The expression of common DEGs (FDR < 0.001) from two RNA sequencing experiments is shown in two heatmaps. The DEGs’ association with selected KEGG pathways is shown in the table on the right. The siRNA used in other panels is named siRNA#1, and the alternative one is siRNA#2.

### Bayesian network reveals central hub genes regulated by CYB5R4

To screen essential regulators downstream of CYB5R4 in an unbiased fashion, we treated HAECs with various concentrations of *CYB5R4* siRNA to observe the correlation between *CYB5R4* and differentially expressed genes (DEGs). To limit the computation to a practical space, only the most significant 230 DEGs (FDR < 0.05) were selected for the analysis (**Fig. 3b**). The candidate genes were used to construct a random directed network which was used to infer the structure of a Bayesian network with gene expression values and a prior probability prior based on the STRING database (**Fig. 3c**). We repeated the learning process to evaluate edge frequencies. Meanwhile, random Bayesian networks were generated by shuffling the gene expression values to serve as the null hypothesis. A consensus network was then constructed by comparing the real networks to random ones and keeping 863 edges with FDR < 0.05 (**Fig. 3d, Supplementary** Fig. 4). In the consensus network, the importance of candidate genes was ranked by their authority score, hub scores, and out-to-in edge ratios (**Fig. 3e**). As a result, three genes, RRM2, TK1, and BIRC5 were determined as the central hub genes.

To rule out potential siRNA off-targets, we transfected HAECs with an alternative siRNA and repeated the RNA sequencing experiment. Compared to the siRNA used for the Bayesian network experiments (siRNA #1), the alternative (siRNA #2) resulted in consistent changes in genes involved in cell cycle and DNA replication (**Fig. 3f**). The three central hub genes are all highly significant in both datasets.

### RRM2 overexpression rescues CYB5R4 knockdown-induced proliferation defects in ECs

Although Bayesian networks can help identify highly informative genes in the gene regulatory network, the true causal relationship between genes must be verified using variable-controlled experiments. Because all central hub genes were downregulated in *CYB5R4*-silenced ECs, we designed a screening method to examine whether restoring the expression of down-regulated central hub genes can rescue the proliferation defects due to the loss of CYB5R4 (**Fig. 4a**). In a pool of HAECs, each carrying a doxycycline-inducible cassette expressing a single gene of interest in the genome, *CYB5R4* was silenced by siRNA transfection. The relative quantification of transgene copies showed that *RRM2* was enriched in the doxycycline-treated samples, suggesting that forcing cells to express RRM2 increased cell proliferation despite the loss of CYB5R4 (**Fig. 4b**). Although the other two central hub genes, *TK1* and *BIRC5*, tended to be positively enriched by doxycycline, the difference was similar to non-central hub genes, such as *CDC6*, *TPX2*, *AARS1, and CYB5R3*, and not statistically significant. In comparison, the cells carrying *EGFP* or only the multiple cloning sites (*MCS*) were negatively enriched in doxycycline-treated samples.

**Fig. 4:**
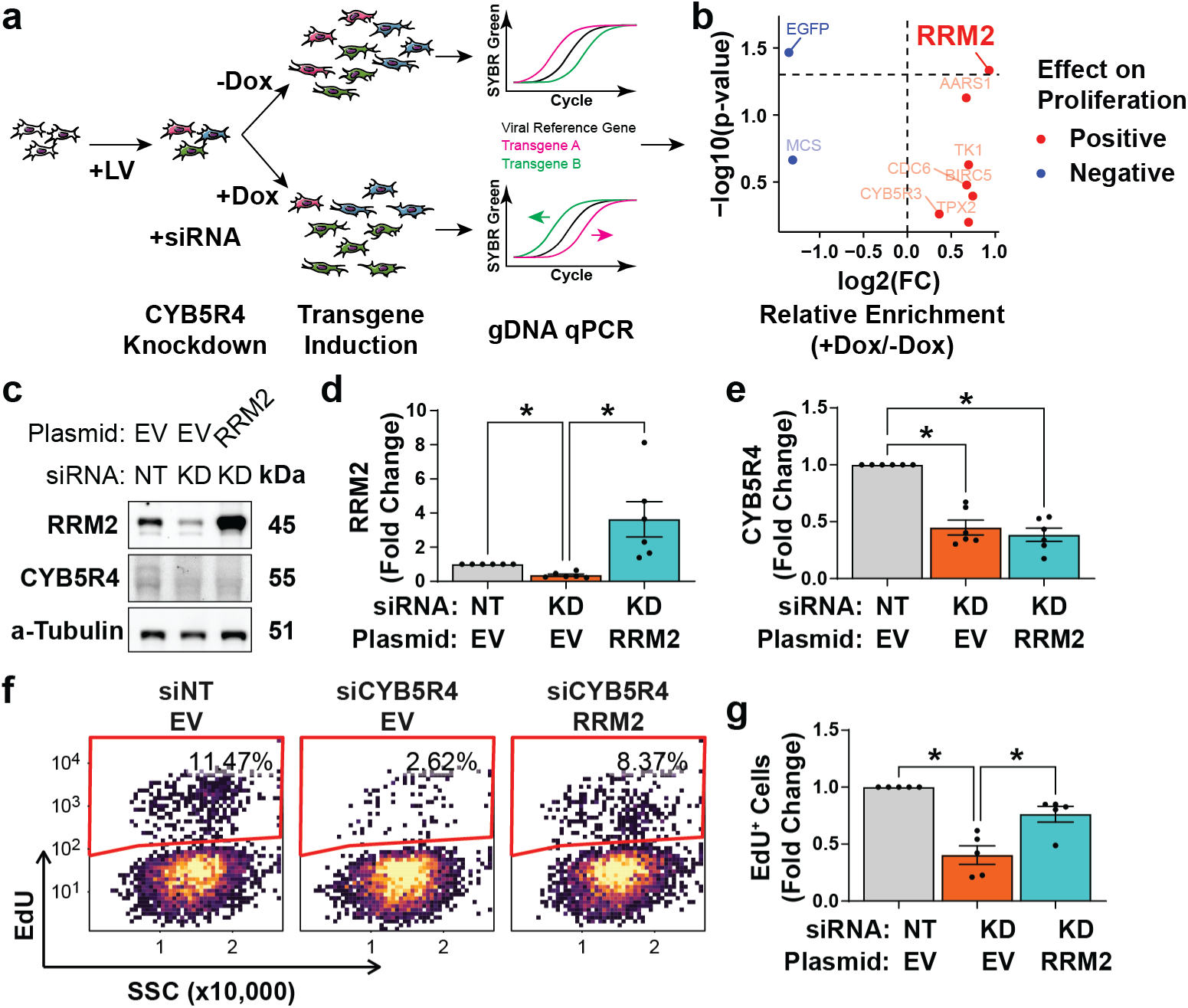
The effect of restoring hub genes in *CYB5R4*-silenced endothelial cells. **(a)** The illustration shows the strategy to screen the effect of hub genes on proliferation. HAECs are transduced with doxycycline (DOX)-inducible lentivirus (LV) expressing hub genes. Cells are pooled and transfected with CYB5R4 siRNA (siCYB5R4). After growing with or without DOX, genome DNA (gDNA)-incorporated transgenes are quantified using quantitative PCR (qPCR). **(b)** The qPCR results are summarized as the relative enrichment of transgenes in DOX-treated cells over the no DOX control. **(c-e)** The representative image and quantification of immunoblotting show CYB5R4 and RRM2 expression with transfection of non-targeting siRNA (NT), CYB5R4-targeting siRNA (KD), empty vector (EV), or RRM2-expressing vector (RRM2). N=6; * indicates p<0.05 using Student’s t-test. **(f, g)** Cell proliferation is examined with EdU incorporation under the same knockdown and overexpression conditions. N=5; * indicates p<0.05 using one-way ANOVA and Tukey’s post-hoc test.

In a separate experiment, HAECs were transfected with siRNA to knock down *CYB5R4*. Consistent with the mRNA expression pattern, the loss of CYB5R4 suppressed RRM2 expression on the protein level (**Fig. 4c-e**). CYB5R4-silenced HAECs were forced to express RRM2 by co-transfection of plasmid DNA expressing *RRM2* (**Fig. 4b**). The EdU incorporation assay confirmed that restoring RRM2 expression improved cell proliferation in *CYB5R4*-knockdown ECs (**Fig. 4f, 4g**).

### CYB5R4 and RRM2 reside in close proximity in ECs

RRM2 is an essential subunit of the ribonucleotide reductase, which converts ribonucleotides to deoxynucleotides in the cytoplasm^37^. As CYB5R4 is also a cytosolic protein^38^, we examined whether CYB5R4 and RRM2 can interact in ECs using two approaches. First, we transduced HAECs with lentivirus to express CYB5R4 with APEXII fused to its C-terminus. The presence of hydrogen peroxide (H_2_O_2_) and biotin-phenol activates APEXII to induce biotinylation of proteins in proximity. RRM2 was identified in the pool of biotinylated proteins, suggesting that CYB5R4 and RRM2 have a close spatial association (**Fig. 5a**).

**Fig. 5:**
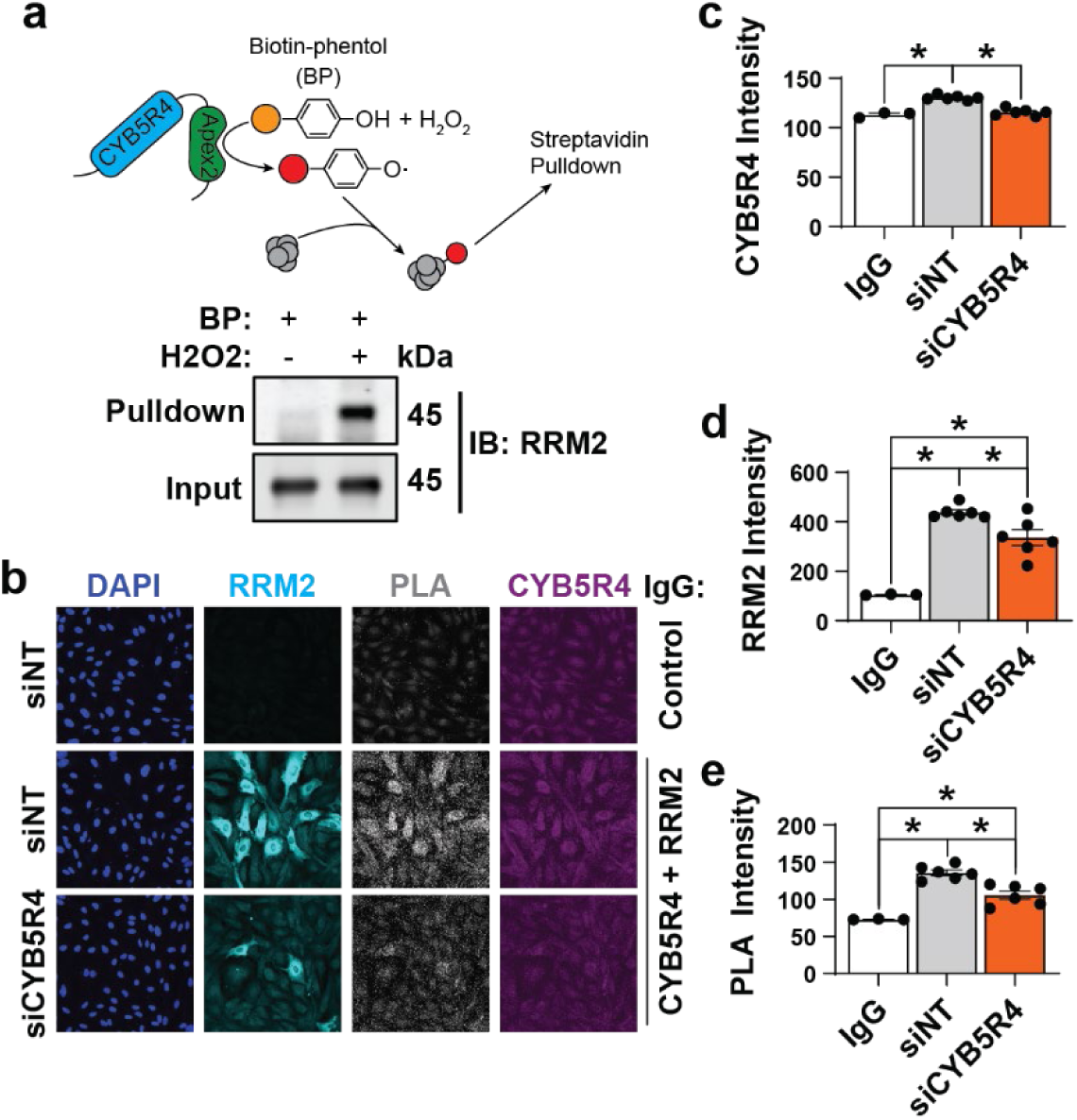
Spatial interaction between CYB5R4 and RRM2. **(a)** The illustration shows APEXII-mediated proximal protein biotinylation. HAECs expressing CYB5R4-APEXII are incubated with biotin phenol and hydrogen peroxide (H_2_O_2_) to induce biotinylation. CYB5R4 is examined in the streptavidin pulldown and input samples using immunoblotting. **(b)** HAECs transfected with non-targeting (siNT) or *CYB5R4*-targeting (siCYB5R4) are used in the proximity ligation assay (PLA). Cells are either incubated with the negative isotype control or RRM2 and CYB5R4-specific antibodies. The RRM2 and CYB5R4 staining or PLA signal intensities are quantified in **(c-e)**. N=3 for the control IgG or 6 for RRM2 and CYB5R4 antibody groups; * indicates p < 0.05 using one-way ANOVA with Tukey’s post-hoc test.

Second, we performed a proximity ligation assay using antibodies against CYB5R4 and RRM2 in HAECs. CYB5R4 siRNA decreased the CYB5R4 and RRM2 immunofluorescent signal, suggesting the antibodies were specific (**Fig. 5b-d**). The proximity ligation signal was only present in cells probed with CYB5R4 and RRM2 antibodies, but not the isotype antibody control, which was decreased by *CYB5R4* silencing (**Fig. 5a, 5e**). The experiment indicated that endogenous CYB5R4 and RRM2 also reside in proximity.

### CYB5R4 regulates the balance of the nucleotide pool

Because CYB5R4 is in the range of physical interaction with RRM2, we investigated whether CYB5R4 regulates ribonucleotide reductase activity. We first performed an unbiased metabolomics screening in siRNA-transfected HAECs using LC-MS. We annotated 775 endogenous compounds with high confidence, among which 49 compounds were annotated as fatty acid or nucleotide metabolites by KEGG (**Fig. 6a**). Monounsaturated fatty acids, such as palmitoleic acid (16:1) and oleic acid (18:1), were both decreased by *CYB5R4* silencing, which was consistent with our isotope-labeled metabolite tracing results (**Fig. 2c**). More compounds involved in nucleotide metabolism showed different levels in cells received *CYB5R4* and non-targeting siRNA.

**Fig. 6:**
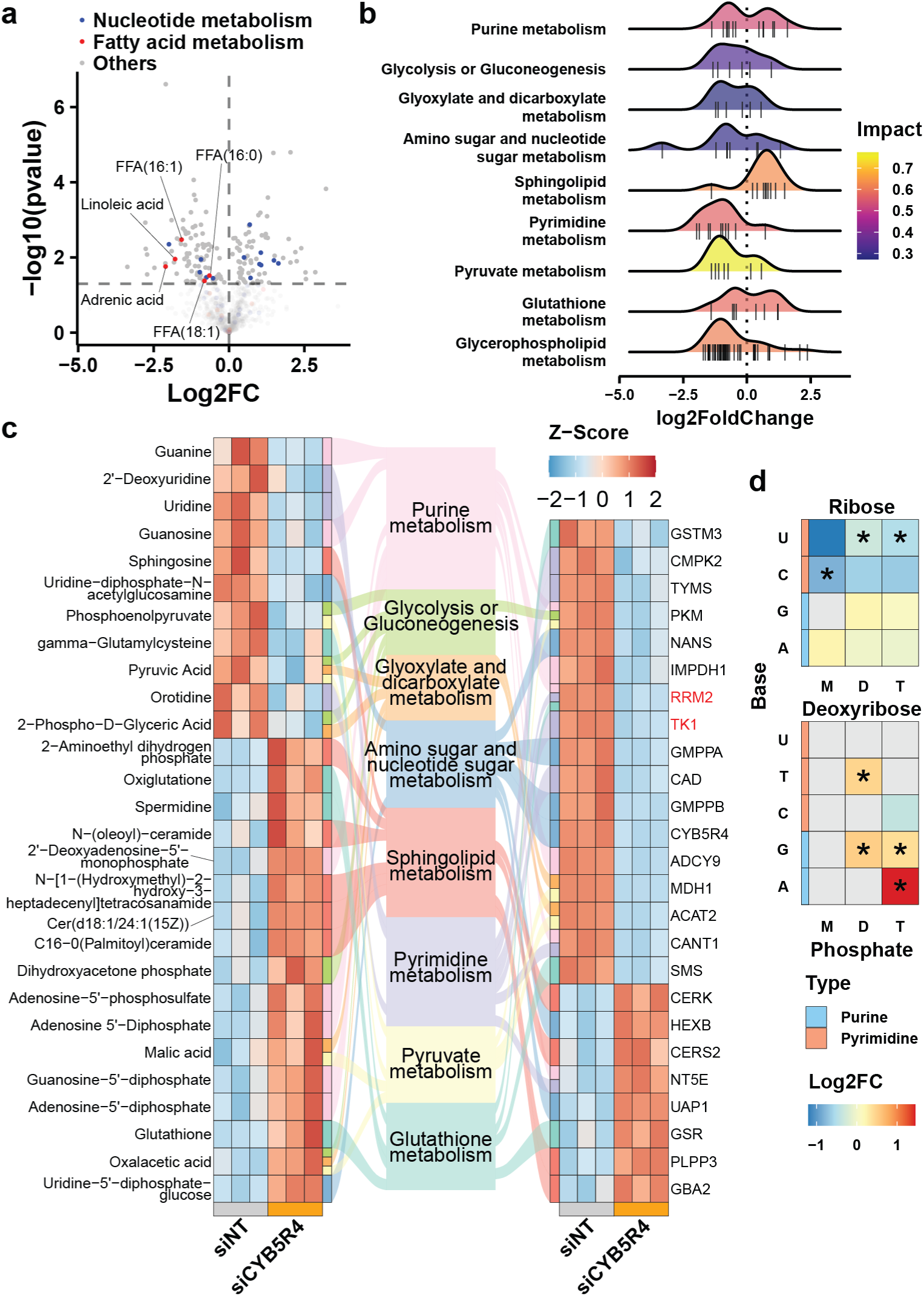
Effect of CYB5R4 on nucleotide metabolism. HAECs transfected with non-targeting or *CYB5R4*-targeting siRNA are used for untargeted metabolomics screening. **(a)** The volcano plot shows significantly up- and down-regulated metabolites (siCYB5R4 over siNT, p < 0.05). Metabolites annotated for nucleotide and fatty acid metabolism in KEGG are highlighted in colors. **(b)** A joint pathway analysis is performed using metabolomics and transcriptomics data. The top 10 significant KEGG pathways are shown in the ridge density plot. The up- and down-regulated metabolites/genes in a pathway are shown as bars below the density plot. **(c)** The connected heatmap shows metabolites/genes involved in the top significant KEGG pathways. The metabolites are shown in the left heatmap, and the genes are shown in the right. The chords connect metabolites and genes to their associated KEGG pathways. **(d)** Nucleotides were measured with a targeted approach in siRNA-transfected HAECs. The heatmap shows relative changes of nucleotides (siCYB5R4 over siNT), classified by the sugar (ribose and deoxyribose), number of phosphate groups [monophosphate (M), diphosphate (D), and triphosphate (T)], and nitrogenous base [uracil (U), thymine (T), cytosine (C), guanine (G), adenine (A)]. Non-detected nucleotides are shown in gray. N = 4; * indicates p < 0.05 using Student’s t-test.

Due to the relatively small coverage of the metabolomics landscape in our screening, a pathway analysis based on the metabolites alone can be problematic. Instead, we performed a joint pathway analysis by combining the metabolomics screening and the RNA sequencing data. The result shows that glycerophospholipids were predominantly decreased in *CYB5R4* knockdown cells (**Fig. 6b and Supplementary** Fig. 5), which can be a result of oxidative stress or proliferation defects. Both purine and pyrimidine metabolism were among the most affected pathways (**Fig. 6b**). While the genes and metabolites involved in purine metabolism did not show a clear direction of changes, they were predominantly down-regulated in pyrimidine metabolism by the loss of CYB5R4 (**Fig. 6b-c**). While both TK1 and RRM2 are involved in pyrimidine metabolism, RRM2 sits at the intersection of purine, pyrimidine, and glutathione metabolism, highlighting its critical position in mediating CYB5R4’s effect in ECs.

To clarify CYB5R4’s regulatory role in nucleotide metabolism, we measured common nucleotide metabolites using a targeted strategy. Among the successfully detected nucleotides, uridine diphosphate (UDP), uridine triphosphate (UTP), and cytidine monophosphate (CMP) were significantly decreased in *CYB5R4* knockdown HAECs. In contrast, deoxythymidine diphosphate (dTDP), deoxyguanosine diphosphate (dGDP), deoxyguanosine triphosphate (dGTP), and deoxyadenosine triphosphate (dATP) were increased by *CYB5R4* silencing (**Fig. 6d**). Overall, *CYB5R4*-silenced ECs show an imbalanced nucleotide pool signified by decreased pyrimidine ribonucleotides.

## Discussion

In this study, we identified CYB5R4 as a critical regulator of angiogenesis and EC proliferation. Although common variants in CYB5R4 do not present a significant risk for PAD in large genome-wide association studies (**Fig. 1a**), the absence of GWAS signals at CYB5R4 may reflect a stringent evolutionary constraint rather than dispensability. Indeed, CYB5R4 shows modest depletion of predicted loss-of-function variation (o/e = 0.58, 90% CI 0.42-0.82) (**Supplementary** Fig. 1). Previously, gene profiling with microarray has shown that CYB5R4 expression declines in aging human atrial tissue^39^, consistent with the known impairment of angiogenesis in older individuals. Interestingly, a genetic variant’s effect on CYB5R4 expression in human skeletal muscle and artery is negatively correlated with the variant’s effect on PAD incidents (**Fig. 1b**). Overall, the human association data suggest that CYB5R4 may have a protective role in the progression of PAD.

In line with these observations, genetic loss of CYB5R4 in mice blunted ischemia-driven angiogenesis. Both global heterozygous and endothelium-specific homozygous CYB5R4 knockout mice exhibited delayed blood flow recovery (**Fig. 1e-f, i-j**). The endothelium-specific CYB5R4 knockout mice diminished capillary proliferation after hindlimb ischemia (**Fig. 1k-m**), underscoring that CYB5R4’s pro-angiogenic function is endothelium-intrinsic. It is worth noting that the global homozygous knockout mice have severe lipoatrophy and early onset diabetes (**Supplementary** Fig. 1), which is consistent with the previous report^20^. The global heterozygous knockouts and the endothelium-specific knockouts showed impaired ischemia-induced angiogenesis, without these systemic metabolic defects. Therefore, decreased expression in human vascular tissue can be a risk factor for PAD independent of lipid metabolic disorder and diabetes. These findings reveal a previously unrecognized role for CYB5R4 in supporting new vessel growth, supporting the protective role of CYB5R4 in PAD.

It is important to note that our global *Cyb5r4* knockout differs from the previously reported *Cyb5r4^tm1Hfb^* line in both allele design and resulting phenotype. The published tm1Hfb allele replaces exon 4, which encodes the heme-binding domain, with a *Pgk-Hyg* expression cassette. Those mice showed no embryonic lethality and developed early insulin-deficient diabetes with progressive lipoatrophy. This strategy may result in exon skipping or splicing to cryptic sites partially rescuing expression^35,40^. Indeed, the author reported mRNA lacking the entire exon 4 from the *Cyb5r4^tm1Hfb^* allele. In comparison, our line deletes exon 3 by CRISPR and exhibits sub-Mendelian viability and lowers body weight around weaning. The more severe phenotype suggests that our line has a more complete loss of gene expression and protein function. Additionally, the *Cyb5r4^tm1Hfb^* mice were generated on the 129S4/SvJae background and mated to BALB/cAnN mice, while our knockout mice were created and mated on the C57BL6/J background, which may also contribute to the phenotypic differences. Nonetheless, both global knockout strains showed consistent defects in systemic lipid metabolism, underscoring the necessity to evaluate EC function with our endothelium-specific tamoxifen-inducible *Cyb5r4* knockout mice.

Mechanistically, our data indicate that CYB5R4 promotes EC proliferation through a pathway independent of its canonical role in fatty acid desaturation. CYB5R4 is known to supply electrons to stearoyl-CoA desaturase (SCD) for the formation of monounsaturated fatty acids^20^. As expected, we observed that CYB5R4 silencing impairs SCD activity in ECs, leading to impaired conversion of stearoyl CoA to oleoyl CoA (**Fig. 2c**). Although the silencing of either SCD or CYB5R4 delayed scratch wound recovery (**Fig. 2b,3**), SCD knockdown paradoxically increased the S-phase fraction (**Fig. 2f-g**). It is possible that the loss of SCD caused cells to be arrested in the late S phase. In comparison, CYB5R4 knockdown caused G1-to-S transition defect (**Fig. 2f-g**). More importantly, supplementing palmitoleic or oleic acids did not rescue proliferation in CYB5R4-deficient cells (**Fig. 2h**). Thus, while CYB5R4 contributes to fatty acid desaturation, its pro-proliferative effect operates via an SCD-independent pathway more directly coupled to cell-cycle control.

Transcriptomic analysis of CYB5R4-deficient ECs reinforced this notion by revealing downregulation of cell-cycle and DNA replication programs. Gene set enrichment analysis showed that CYB5R4 knockdown suppresses many S-phase-associated genes (e.g., DNA polymerase subunits, cyclins, mitotic regulators) while upregulating oxidative stress-related genes (**Fig. 3a, Supplementary** Fig.3). Notably, core cell-cycle regulators were broadly diminished in the absence of CYB5R4, correlating with the observed G1-S checkpoint failure (**Fig. 3a,f**). These transcriptomic changes are consistent with our phenotypic data, suggesting that CYB5R4 is required to maintain the pro-proliferative transcriptional program of ECs.

Our integrated transcriptomic and metabolomic analyses converged on nucleotide metabolism as a major pathway regulated by CYB5R4. Among the most striking transcript-level changes in CYB5R4-deficient ECs was the suppression of RRM2, the gene encoding the regulatory subunit of ribonucleotide reductase (**Fig. 3e,f**), which was confirmed on the protein level (**Fig. 4c,g, 5b,d**). RRM2 emerged as a central hub gene in our Bayesian network inference of CYB5R4’s downstream targets, alongside other cell-cycle genes (*TK1* and *BIRC5*). We acknowledge that Bayesian network modeling is an inferential tool with inherent limitations, as it infers putative causal relationships from correlation patterns and is sensitive to confounding variables, which has led to debatable reliability in establishing true causation. In our study, we used the Bayesian network approach exclusively as an unbiased screen to generate hypotheses, not as definitive proof. To strengthen confidence in the network’s predictions, we performed rigorous experimental validation (**Fig. 4**). Notably, *RRM2* stood out as a consistently and significantly downregulated gene in CYB5R4-silenced cells with two independent siRNAs (**Fig. 3f**). Restoring RRM2 levels in CYB5R4-deficient ECs partially rescued their proliferation defect, whereas parallel overexpression of other putative hubs (such as *TK1* or *BIRC5*) did not produce a significant rescue (**Fig. 4**). This genetic rescue experiment establishes *RRM2* as a key effector downstream of CYB5R4, lending credence to the causal chain suggested by the network model.

Interestingly, we also found evidence of a physical proximity between CYB5R4 and RRM2 proteins inside ECs. Using peroxidase-mediated protein labeling and proximity ligation assays, we detected CYB5R4 and RRM2 in close spatial association in the cytosol of ECs (**Fig. 5**). This result suggests that CYB5R4 might interact with the ribonucleotide reductase complex or an associated protein complex. However, the molecular nature and significance of this proximity remain unclear. One speculation is that CYB5R4 could influence RRM2 or RNR activity through a redox-based mechanism. RRM2 contains a di-iron cofactor that generates a stable tyrosyl radical essential for RNR catalytic function^41^. The tyrosyl radical in mammalian RRM2 is not stable, but it is unclear how the diiron center is recycled to regenerate the tyrosyl radical^42^. Interestingly, CYB5R4’s known substrate SCD also has a di-iron center, which is essential for its activity^43^. It is possible that CYB5R4 reduces the di-iron center in RRM2 and maintains RRM2’s tyrosyl radical, which is a subject of future investigation.

Classically, the RNR enzyme’s activity is maintained by thioredoxin/thioredoxin reductase, which regenerates the active site cysteine residues, and by tight allosteric regulation^44^. It is intriguing to consider that CYB5R4 may have the capacity to modulate the redox state of RRM2 or augment electron transfer in nucleotide reduction. At this stage, however, we have no direct evidence that CYB5R4 enzymatically influences RNR activity, and other interpretations are equally plausible.

For instance, the loss of CYB5R4 could indirectly downregulate RRM2 via cell-cycle signaling pathways (e.g., through reduced E2F transcriptional activity in G1-arrested cells)^45,46^, in which case the CYB5R4-RRM2 proximity may be related to protein complex assembly rather than a direct enzymatic interaction.

Our metabolomic findings provide additional context but also underscore the complexity of this regulation. CYB5R4 knockdown caused measurable shifts in nucleotide pools: we observed significant decreases in pyrimidine ribonucleotides, such as UDP, UTP, and CMP, and concomitant increases in certain deoxynucleotides such as dTDP, dGTP, and dATP, among the subset of nucleotides reliably detected (**Fig. 6d**). This pattern indicates an imbalance in nucleotide production consistent with RNR dysregulation. For example, accumulation of some dNTPs alongside depletion of others could indicate aberrant feedback control of RNR or an altered flux through pyrimidine vs. purine synthesis pathways. Indeed, joint pathway analysis of our metabolomic and transcriptomic data indicated that pyrimidine metabolism was broadly downregulated in CYB5R4-deficient cells, whereas purine metabolites were less consistently changed or even elevated (**Fig. 6b,c**). RRM2 sits at the intersection of purine and pyrimidine deoxynucleotide production, so an RRM2 defect could certainly disrupt this balance, which inhibits DNA synthesis and cell proliferation^47,48^.

However, we caution that these metabolite changes do not permit definitive conclusions about RNR enzymatic activity. A limitation is that our untargeted metabolomic screen and targeted nucleotide measurement did not have full coverage of the nucleotide pool. Many nucleotide species, particularly ribonucleotide diphosphates (NDPs) and deoxynucleoside diphosphates (dNDPs), were not detected due to technical constraints. Notably, the immediate substrates and products of RRM2, dADP, dGDP, dCDP, and dUDP, were largely below detection in our analysis, leaving us inferential evidence rather than a complete picture. It is also possible that the observed dNTP accumulations reflect downstream metabolic or cell-cycle effects (e.g., stalled DNA replication leading to feedback accumulation of certain dNTPs) rather than a direct increase in RNR catalytic turnover. Nonetheless, while our data strongly implicate RRM2 as a downstream mediator of CYB5R4, the exact impact of CYB5R4 on RNR enzyme function remains to be clarified.

In conclusion, this study uncovers a previously unappreciated link between CYB5R4 and the nucleotide metabolic machinery that drives angiogenesis. CYB5R4 serves as a bridge connecting redox homeostasis, lipid metabolism, and the cell cycle in ECs. By ensuring adequate and balanced dNTP synthesis through RRM2, CYB5R4 enables ECs to proliferate and form new blood vessels under ischemic conditions. These findings not only shed light on the fundamental biology of angiogenesis but also suggest that metabolic reinforcement of ECs, via the CYB5R4-RRM2 axis, could be a novel angle to improve therapeutic angiogenesis in diseases like PAD.

## Acknowledgements

We thank the University of Pittsburgh Innovative Technologies Development Core and Dr. Sebastien Gingras for the generation of CYB5R4 global and conditional knockout mice using CRISPR/Cas9 technology. We acknowledge Dr. Xinghua Lu for the advice on causal inference with RNA sequencing data and Dr. Marco Scutari for the instructions on using bnlearn for Bayesian network analysis.

The Genotype-Tissue Expression (GTEx) Project was supported by the Common Fund of the Office of the Director of the National Institutes of Health, and by NCI, NHGRI, NHLBI, NIDA, NIMH, and NINDS. The data used for the analyses described in this manuscript were obtained from: dbGaP accession number phs000424.v8.p2 on 09/25/2024.

## Sources of Funding

This work was supported by the National Institutes of Health (NIH) grants: NIH/NHLBI R35HL161177 (A.C.S.), R01GM125944 and R01AR084311 (F.J.S.), 1F31 HL172595 (N.C.).

Work performed in the Health Sciences Mass Spectrometry Core (RRID:SCR_025222) and services and instruments used in this project were graciously supported, in part, by the University of Pittsburgh and the Office of the Senior Vice Chancellor for Health Sciences and NIH S10OD032141 (S.L.G.).

## Author Contributions

S.Y. conceived the study hypothesis, designed and performed the majority of experiments, analyzed data, and wrote the manuscript. S.A.H. performed the animal surgery and maintained the animal colony. N.C.C. and F.J.S. designed and performed experiments to measure SCD activity and analyzed the data. S.J.M. and S.L.G. designed and performed experiments for targeted nucleotide measurements and analyzed the data. A.C.S. conceived the study hypothesis, supervised the study, and wrote the manuscript. All authors reviewed and edited the manuscript.

---

**Fig. S1:**
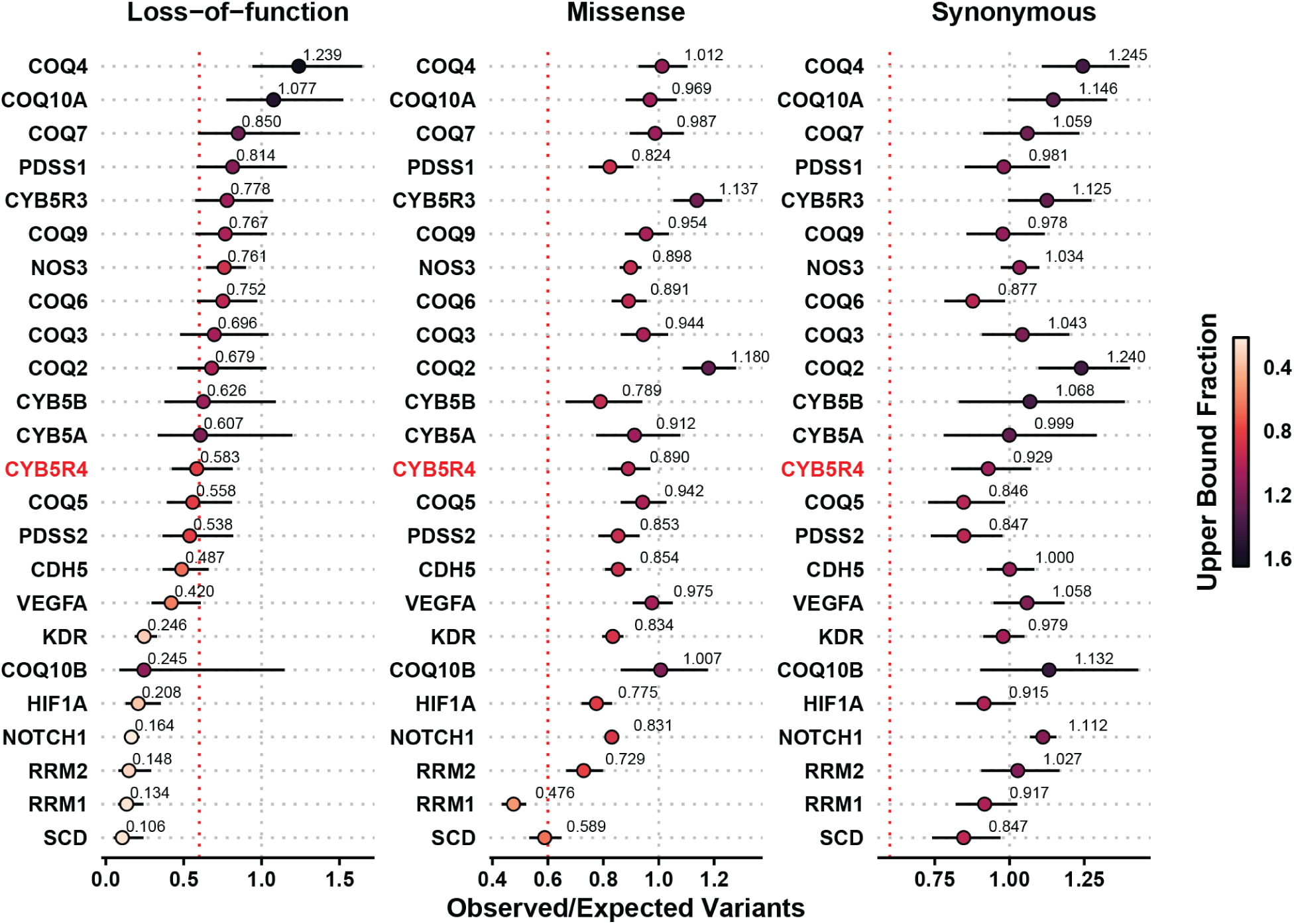
The gnomAD gene constraint metrics for loss-of-function, missense, and synonymous genetic variants. The gene constraint data for CYB5R4 and selected genes are retrieved from the gnomAD. The dot position indicates the ratio of the observed variants to the expected variants, while the range line shows the lower and upper bound fractions. The color of the dots is mapped to the observed/expected upper bound fraction. The red vertical dashed line indicates the gnomAD recommended threshold for loss-of-function observed/expected upper bound fraction (LOEUF=0.6).

**Fig. S2:**
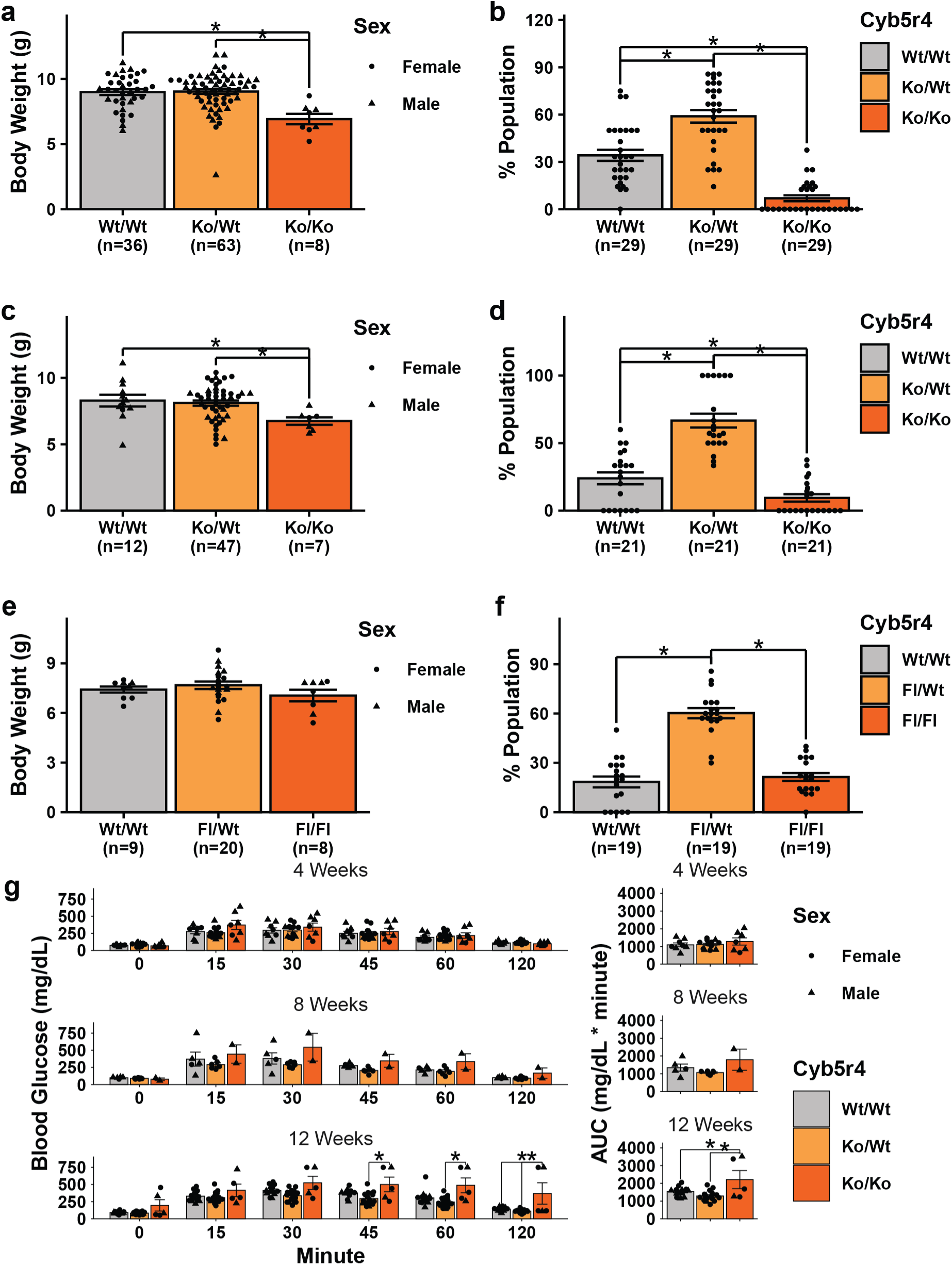
Characterization of *CYB5R4* knockout strains. (a-f) Body weight measurements of 21-day-old pups and the genotype ratio for different *CYB5R4* knockout strains. **(a-b)** 16 litters from 5 global CYB5R4^Ko/Wt^ x CYB5R4^Ko/Wt^ breeders. N values are labeled in the figure; * indicates p < 0.05 using one-way ANOVA with Tukey’s post-hoc test. **(c-d)** 21 litters from 7 global CYB5R4^Ko/Wt^ x CYB5R4^Ko/Wt^ breeders derived from the CYB5R4-floxed strain. * indicates p < 0.05 using one-way ANOVA with Tukey’s post-hoc test. **(e-f)** 5 litters from 3 CYB5R4^Fl/Wt^ x CYB5R4^Fl/Wt^ breeders (endothelium-specific tamoxifen-inducible strain). * indicates p < 0.05 using one-way ANOVA with Tukey’s post-hoc test. **(g)** Intraperitoneal glucose tolerance test combined global CYB5R4 knockout mice combining the original knockout strain and CYB5R4-flox allele converted global knockout mice.

**Fig. S3:**
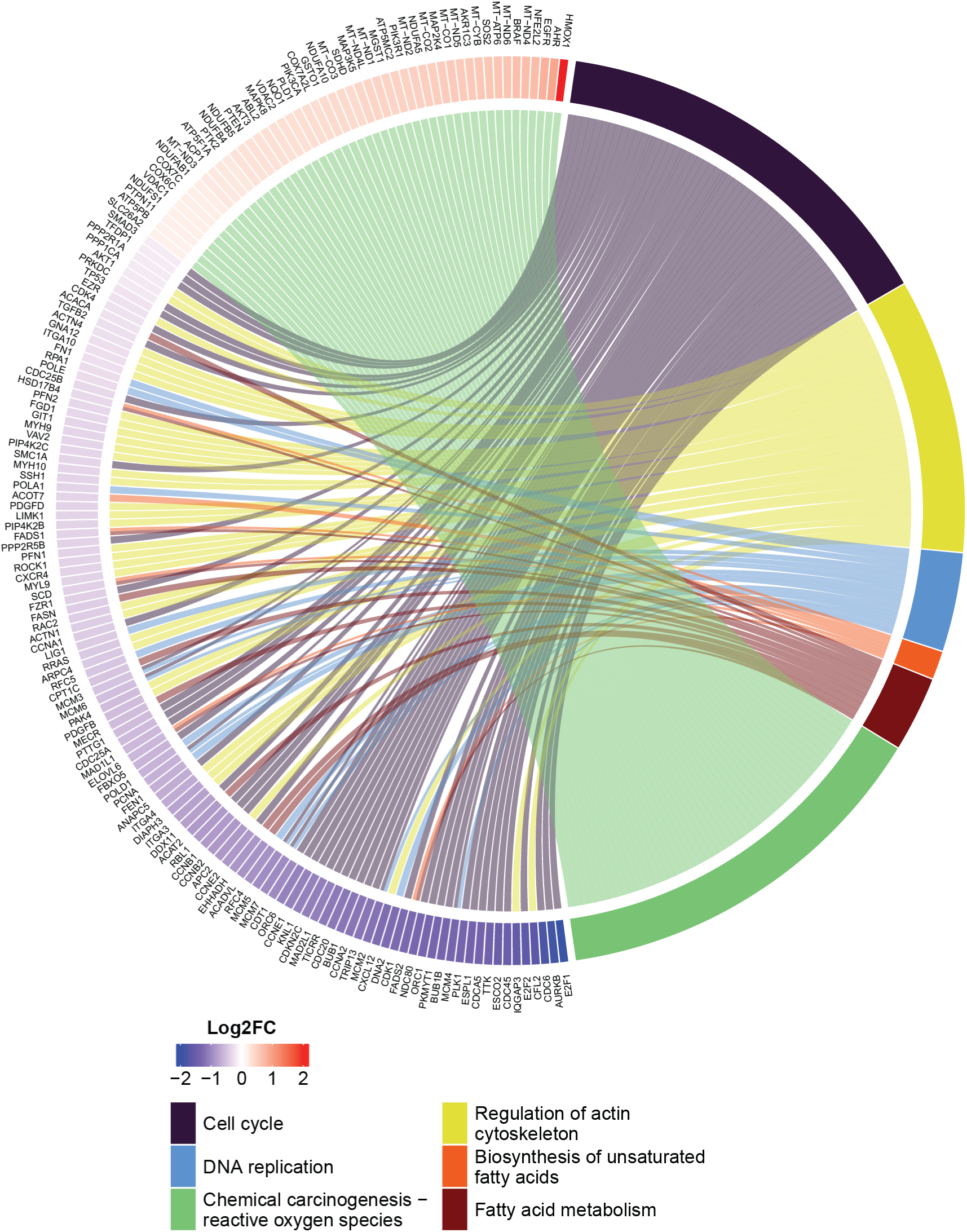
Chord plot for selected pathways from gene set enrichment analysis. Human aortic endothelial cells (HAECs) are transfected with 10 μM non-targeting (siNT) or 10 μM of CYB5R4-targeting (siCYB5R4) siRNA for RNA sequencing. Core enrichment genes are linked to the selected significant (FDR < 0.05) Kyoto Encyclopedia of Genes and Genomes (KEGG) pathways from a gene set enrichment analysis.

**Fig. S4:**
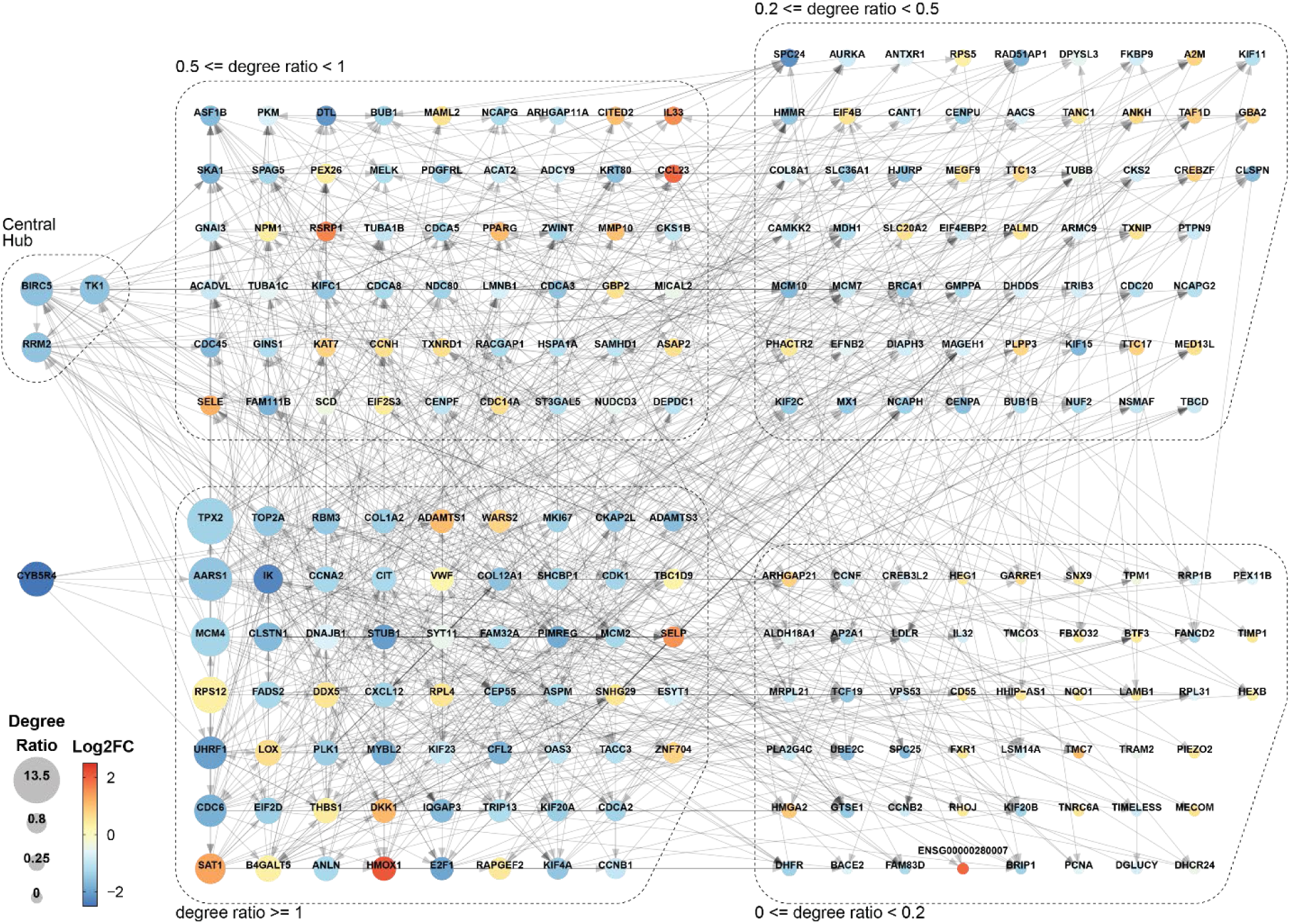
Complete consensus network. Significant edges from Bayesian networks are summarized in a consensus network. Genes are grouped by their out-to-in degree ratios.

**Fig. S5:**
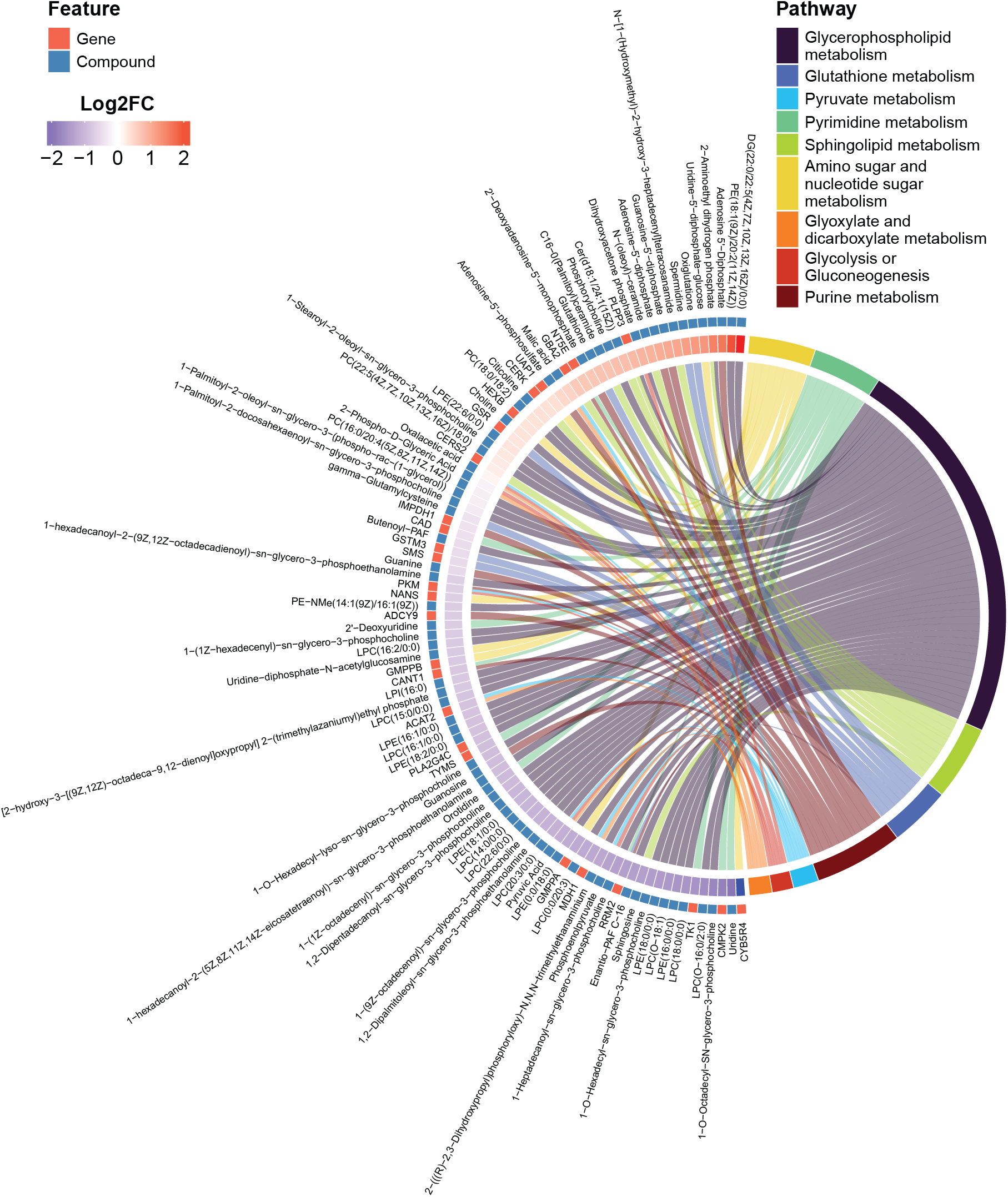
Chord plot for all top pathways from the joint pathway analysis. All top 10 significant pathways are included. Genes or compounds are linked to the associated pathways.

## Graphical Summary

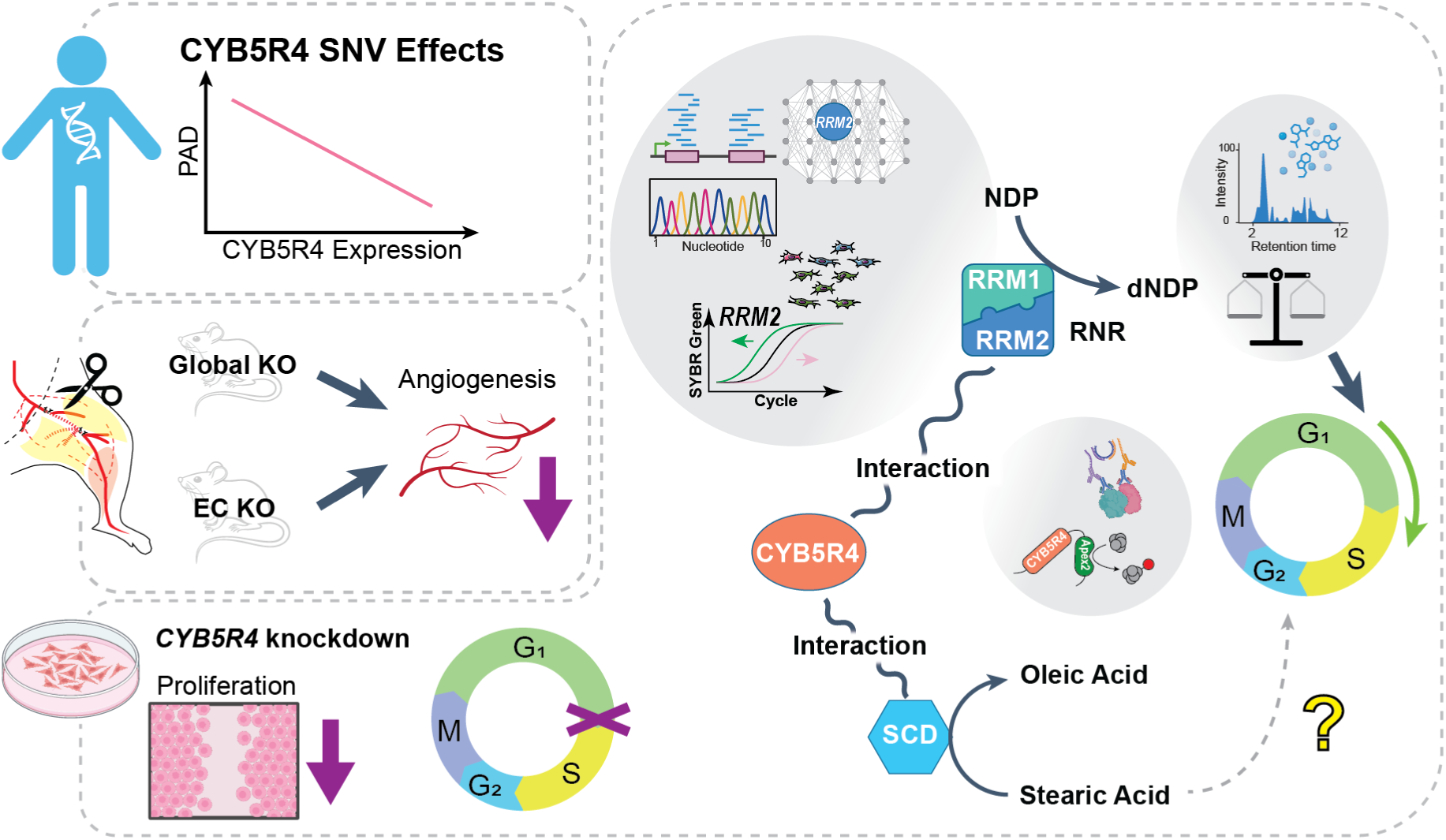

**Supplementary Table 1.**
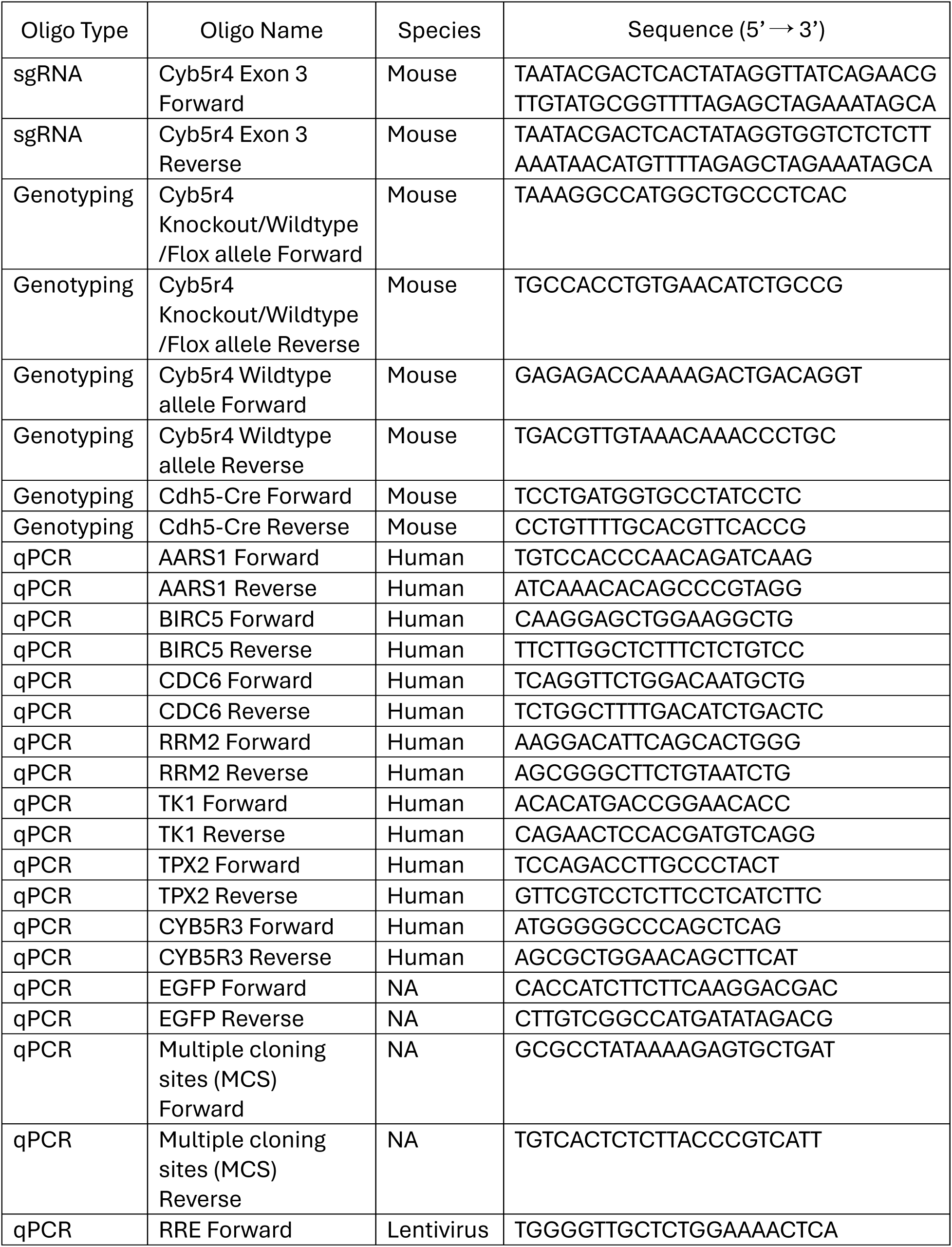
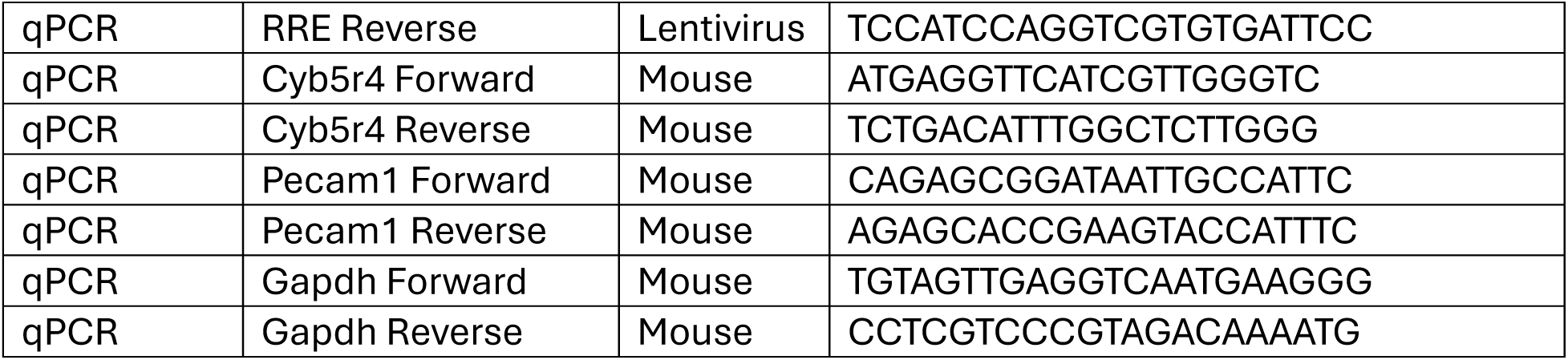

